# Human mutations in *SLITRK3* implicated in GABAergic synapse development in mice

**DOI:** 10.1101/2022.12.19.520993

**Authors:** Stephanie Efthymiou, Wenyan Han, Muhammad Ilyas, Jun Li, Yichao Yu, Marcello Scala, Nancy T. Malintan, Muhammad Ilyas, Nikoleta Vavouraki, Kshitij Mankad, Reza Maroofian, Clarissa Rocca, Vincenzo Salpietro, Shenela Lakhani, Eric J. Mallack, Timothy Blake Palculict, Hong Li, Guojun Zhang, Faisal Zafar, Nuzhat Rana, Noriko Takashima, Hayato Matsunaga, Queen Square Genomics, SYNAPS Study Group, Pasquale Striano, Mark F. Lythgoe, Jun Aruga, Wei Lu, Henry Houlden

**Affiliations:** Department of Neuromuscular disorders, UCL Queen Square Institute of Neurology, London WC1N 3BG, UK; Synapse and Neural Circuit Research Unit, National Institute of Neurological Disorders and Stroke, National Institutes of Health, Bethesda, MD 20892, USA; IIUI, Department of Bioinformatics & Biotechnology, Islamabad, 44000, Pakistan; Centre for Advanced Biomedical Imaging, Division of Medicine, University College London, 72 Huntley Street, London, WC1E 6DD, UK; Center for Neurogenetics, Feil Family Brain and Mind Research Institute, Weill Cornell Medicine, New York, NY, USA; GeneDx, Gaithersburg, MD, USA; Emory University School of Medicine, Department of Pediatrics, Division of Neurology, Atlanta, GA, USA; Department of Pediatric Neurology, Children’s Healthcare of Atlanta, Atlanta, GA, USA; Department of Pediatrics, Multan Hospital, Multan, Pakistan; Laboratory for Behavioral and Developmental Disorders, RIKEN Brain Science Institute (BSI), Wako-shi, Saitama 351-0198, Japan; Department of Medical Pharmacology, Nagasaki University Institute of Biomedical Sciences, Nagasaki 852-8523, Japan; Department of Neurosciences, Rehabilitation, Ophthalmology, Genetics, Maternal and Child Health, Università Degli Studi di Genova, Genoa, Italy; Pediatric Neurology and Muscular Diseases Unit, IRCCS Istituto Giannina Gaslini, Genoa, Italy; Centre for Omic Sciences, Islamia College Peshawar, Peshawar, Pakistan; School of Pharmacy, University of Reading, Reading, RG6 6AX, UK; Department of Mathematics and Statistics, University of Reading, Reading, RG6 6AX, UK; Department of Radiology, Great Ormond Street Hospital, London, United Kingdom; Developmental Neurosciences Department, UCL Great Ormond Street Institute of Child Health, London, United Kingdom

**Author notes:** **Corresponding author:** Henry Houlden, MD, PhD;, Jun Aruga, M.D. PhD,; Wei Lu, PhD.

**Keywords:** Slitrk3, GABAergic synapse development, epilepsy, global developmental delay

## Abstract

We report on biallelic homozygous and monoallelic *de-novo* variants in *SLITRK3* in 3 unrelated families presenting with epileptic encephalopathy associated with a broad neurological involvement characterized by microcephaly, intellectual disability, seizures, and global developmental delay. *SLITRK3* encodes for a transmembrane protein that is involved in controlling neurite outgrowth and inhibitory synapse development and that has an important role in brain function and neurological diseases. Using primary cultures of hippocampal neurons carrying patients’ SLITRK3 variants and in combination with electrophysiology, we demonstrate that recessive variants are loss-of-function alleles. By analyzing the development and phenotype of SLITRK3 KO (*SLITRK3^-/-^*) mice, we bring additional evidence of enhanced susceptibility to pentylenetetrazole-induced seizure with the appearance of spontaneous epileptiform EEG, as well as developmental deficits such as higher motor activities and reduced parvalbumin interneurons. Taken together, our results exhibit impaired development of peripheral and central nervous system and support a conserved role of this transmembrane protein in neurological function. Our study delineates an emerging spectrum of human core synaptopathies caused by variants in genes that encode SLITRK proteins and essential regulatory components of the synaptic machinery. The hallmark of these disorders is impaired postsynaptic neurotransmission at nerve terminals; an impaired neurotransmission resulting in a wide array of (often overlapping) clinical features, including neurodevelopmental impairment, weakness, seizures, and abnormal movements. The genetic synaptopathy caused by SLITRK3 mutations highlights the key roles of this gene in human brain development and function.

## Introduction

Synapse development is a multi-step process coordinated by various molecules acting in a highly spatially and temporally controlled manner (1). Synaptic cell adhesion molecules (CAMs) are an example of such molecules essential for the establishment and maturation of synaptic connections. Recent studies have identified a growing number of synaptic CAMs such as slit and NTRK-like family proteins (Slitrks) that are single-passing transmembrane proteins and bind to presynaptic protein tyrosine phosphatase (2–4). Members of the Slitrk family contain twelve N-terminal leucine-rich repeat (LRR) domains and C-terminal regions that share homology with neurotrophin receptors (5). They are expressed predominantly in neural tissues of the central nervous system and have neurite-modulating activity (6, 7). Even though such molecules are involved in various stages of synapse development, their function diversity still remains largely unclear.

Recent studies in humans and genetic mouse models have led to the identification of Slitrk family member genes as candidate genes that may be involved in the development of neuropsychiatric conditions such as obsessive-compulsive spectrum disorders, schizophrenia, Tourette syndrome, or trichotillomania (8, 9). Additionally, Slitrk3 abnormality has also been strongly associated with multiple cancers, such as human epithelial ovarian cancer (10) and squamous cell lung cancer (11).

SLITRK3 (Slit and Trk-like family member 3), is a synaptic cell adhesion molecule highly expressed at inhibitory synapses and enhances inhibitory synapse formation (12) (4). Recent studies have provided additional evidence to our understanding of the mechanisms of GABAergic synapse formation by revealing a direct extracellular protein-protein interaction between SLITRK3 and another inhibitory postsynaptic cell-adhesion molecule, Neuroligin 2 (NL2) (13) and a critical role of the SLITRK3-gephyrin interaction in stabilization of pre- and post-synaptic compartments during development (14). Collectively, these studies have shown that SLITRK3, through its extracellular domain, selectively regulates inhibitory synapse development via the trans-synaptic interaction with presynaptic cell adhesion molecule, receptor protein tyrosine phosphatase δ (PTPδ), the *cis* interaction with postsynaptic NL2, and the intracellular interaction with gephyrin. In this way, they exert differential, sometimes cooperative effects on GABAergic synapse formation depending on the developmental stage of the system. Furthermore, other recent studies have shown a critical role of a conserved tyrosine residue, Y969, in SLITRK3 carboxyl-terminus in GABAergic synapse development (14, 15), indicating SLITRK3 may mediate signaling through both extracellular and intracellular domains.

In the brain, many types of interneurons make functionally diverse inhibitory synapses onto principal neurons. In the mammalian brain GABAergic synapses provide an important level of inhibitory balance to glutamatergic excitatory drive, therefore, controlling neuronal excitability and synaptic plasticity. As GABAergic inhibition is important in almost every aspect of brain physiology, and the dysregulation of GABAergic synapse development has been implicated in neurological and neuropsychiatric disorders (16–19), it is critical to understand the molecular determinants of GABAergic synapse formation. Prevention and treatment of brain disorders will also depend partly on restoration of GABAergic function and/or inhibitory/excitatory balance.

In this study, we report on biallelic and monoallelic variants in *SLITRK3* in 3 families presenting with epileptic encephalopathy associated with neurodevelopmental findings that include seizures, motor delay, microcephaly, hyperactivity, and MRI brain abnormalities. Using primary cultures of hippocampal neurons carrying patients’ SLITRK3 variants and in combination with electrophysiology, we characterized the functional effects of these variants on inhibitory synapses. Furthermore, by analyzing the development and phenotype of SLITRK3 KO (*SLITRK3^-/-^*) mice, we looked at survival, seizure susceptibility, and any developmental deficits to correlate to the patients’ phenotypes.

## Materials and Methods

After local institutional review board approval of this study and informed consent from the families, we collected blood samples from the three patients and their parents and extracted DNA using standard procedures. Other families were identified by screening genomic data sets from several diagnostic and research genetic laboratories internationally, as well as using GeneMatcher (20).

### Exome sequencing

To investigate the genetic cause of the disease, WES was performed in the three affected siblings (Fig.1A, II-1, II-2 and II-3) as described earlier and analysis was carried out to fit a recessive model (i.e., homozygous or compound heterozygous), and/or located in genes previously associated epilepsy or other neurological phenotypes. The candidate variants were confirmed after filtering and interpretation according to the American College of Medical Genetics and Genomics/Association for Molecular Pathology guidelines (21) and segregation analyses were carried out using Sanger sequencing. Homozygosity mapping was performed on the genetic data from the 3 affected individuals and unrelated healthy controls age- and ethnically matched (22). Genomic DNA from the submitted specimens analysed at GeneDx was enriched for the complete coding regions and splice site junctions for most genes of the human genome using a proprietary capture system developed by GeneDx for NGS with CNV calling (NGS-CNV). Using a custom-developed analysis tool (XomeAnalyzer), data were filtered and analysed to identify sequence variants and most deletions and duplications involving three or more coding exons.

**Fig 1.**
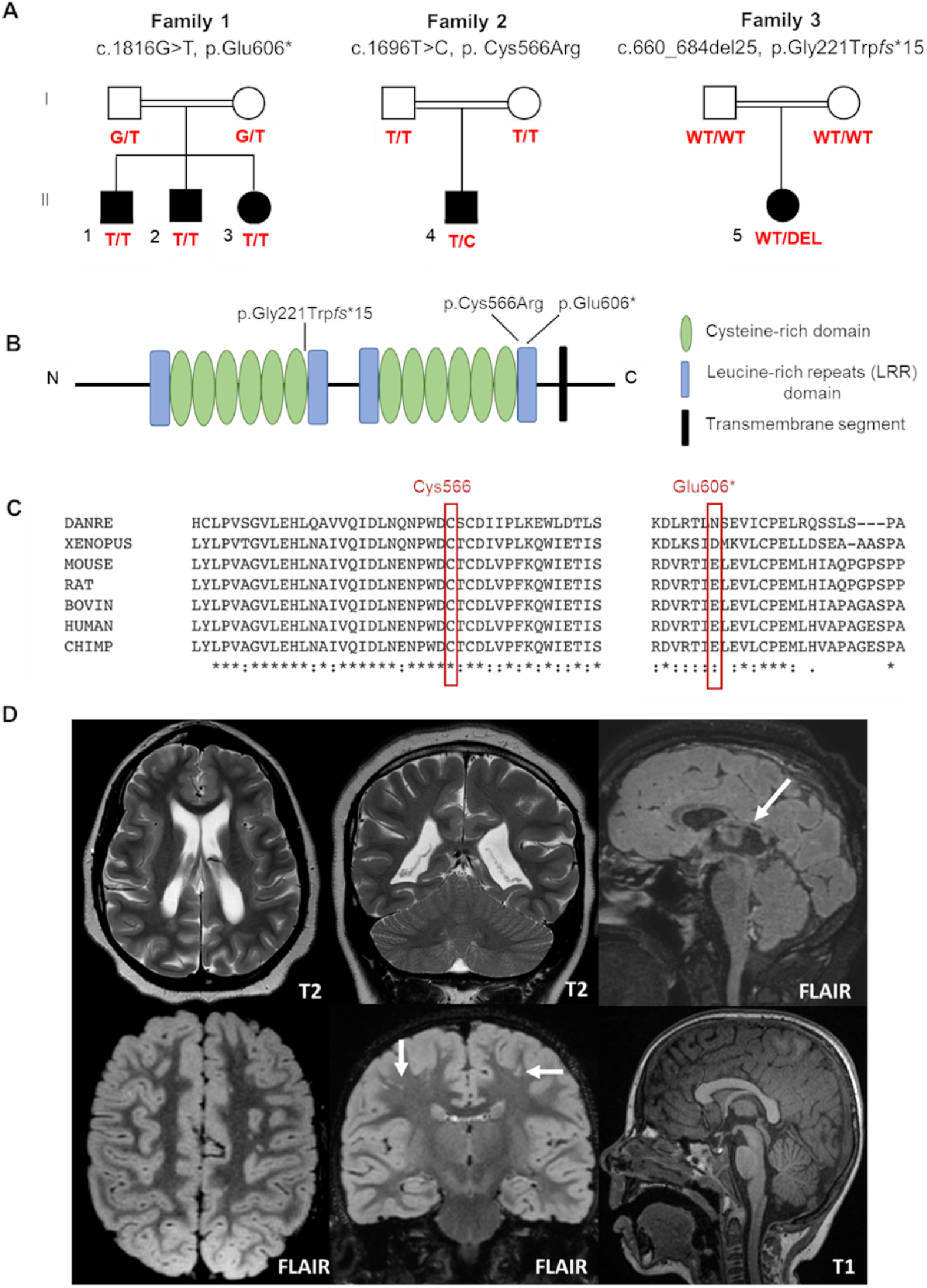
Pedigrees and genetic findings of the families carrying biallelic *SLITRK3* variants. (A) Pedigrees and segregation results (+ represents the presence of the variant) of the three families (B) A schematic representation of the SLITRK3 protein showing the position of all variants identified. (C) Overview of the whole regions of homozygosity (ROH) in the exome of each affected individual in family 1. The region of homozygosity surrounding the SLITRK3 variant is indicated in red bracket on the left. (D) Inter-species alignment performed with Clustal Omega shows the complete conservation down to invertebrates of the amino acid residues affected by the substitutions. (E) top panel (patient 5): Note is made of loss of white matter in both cerebral hemispheres with scalloping of the ventricular margins. On the sagittal FLAIR sequence, thinning of the posterior aspect of the corpus callosum is noted (arrow). Appearances would be compatible with periventricular leukomalacia in the context of white matter injury of prematurity, below panel (patient 4): Multiple tiny foci of white matter signal abnormality are observed in the subcortical white matter of both cerebral hemispheres. This is shown best on the coronal FLAIR image (arrows). The midline sagittal T1 is normal.

### Molecular cloning

SLITRK3 constructs were cDNA synthesised by Genescript (USA) into pCMV-3Tag-4A by replacing the CDs for these inserts via using restriction enzymes BamHI and XhoI (NEB), giving rise to all SLITRK3 plasmid variants. The variants generated were pCMV_SLITRK3[WT], pCMV_SLITRK3[E6O6X] and pCMV_SLITRK3[C566R]. Three Myc epitopes were then inserted before the stop codon into the three plasmids with the QuickChange Site-Directed Mutagenesis Kit (Agilent Technologies) to get SLITRK3-WT-Myc, SLITRK3-C566R-Myc and SLITRK3-E606X-Myc plasmids.

### Western blotting

For plasmid validation, SLITRK3-WT-Myc, SLITRK3-C566R-Myc, or SLITRK3-E606X-Myc were transfected in HEK293T cells using a calcium phosphate transfection reagent. Protein concentration in the soluble supernatant was quantified with the standard BCA method. An equal amount of loading samples was mixed with an equal volume of 2x SDS loading buffer and boiled for 5 min at 95°C, and then separated using pre-casted 10% SDS-PAGE gels (BioRad). The proteins were transferred onto PVDF membranes, blocked, and incubated with primary antibodies as: rabbit polyclonal anti-SLITRK3, (Proteintech, Cat#: 21649-1-AP), recognizing SLITRK3 C-terminus, rabbit monoclonal anti-Myc ((71D10), Cell Signaling Technology, Cat#: 2278), and mouse anti-a-Tubulin (Sigma, Cat#: T8203) overnight at 4°C. The PVDF membranes were then washed three times with 0.1% TBST and incubated with HRP-conjugated secondary antibodies for 1 hr at RT. Protein was detected with the standard enhanced chemiluminescence (ECL) method.

### Dissociated hippocampal neuronal culture

Dissociated cultures of hippocampal neurons were prepared as described previously (13, 23, 24). Briefly, timed-pregnant mice at E17.5-18.5 were anesthetized on ice and decapitated. The fetal hippocampi were quickly dissected out in ice-cold Hank’s balanced salt solution, triturated with a sterile tweezer, and digested with papain (Worthington, LK003176) solution at 37°C for 30 min. After 5 min centrifuging at 800 rpm. at room temperature (RT), the pellet was resuspended in DNaseI containing Hank’s solution and then was mechanically dissociated into single cells by gentle pipetting up and down. Cells were then transferred into Hank’s solution mixed with trypsin inhibitor (10 mg/ml, Sigma, T9253) and BSA (10 mg/ml, Sigma, A9647), and centrifuged at 800 rpm. for 10 min. The pellet was resuspended in neurobasal plating media with 2% fetal bovine serum (FBS) (Gibco, 10437-028), 2% B27 supplements, and L-glutamine (2 mM). Neurons were plated at a density of ~0.9 x 10^5^ cells/well on poly-D-lysine (Sigma, P6407) pre-coated glass coverslips (12 mm) residing in 24-well plates for electrophysiological recording and immunocytochemistry. Culture media were changed by half volume with neurobasal maintenance media containing 2% B27 (GIBCO, 17504-044) supplements and L-glutamine (2 mM) once a week.

### Electrophysiology in neurons

For the miniature inhibitory postsynaptic currents (mIPSC) recording, SLITRK3-WT-Myc, SLITRK3-E606X-Myc, or SLITRK3-C566R-Myc together with pCAGGS-IRES-GFP (SLITRK3:GFP=9:1) were transfected into cultured hippocampal neurons at DIV 14-15 using Lipofectamine 3000 (ThermoFisher, L3000015). 48 hours after transfection, coverslips were transferred to a submersion chamber on an upright Olympus microscope and perfused with aCSF solution supplemented with TTX (0.5 μM), DNQX (20 μM) in an external solution (in mM): 140 NaCl, 5 KCl, 2 CaCl_2_, 1 MgCl_2_, 10 HEPES and 10 D-glucose, adjusted to pH 7.4 with NaOH with 5% CO_2_. GFP fluorescent positive and negative neurons were identified by epifluorescence microscopy. Neurons were voltage-clamped at −70 mV for the detection of mIPSC events. The internal solution for mIPSC recording (in mM): CsMeSO_4_ 70, CsCl 70, NaCl 8, EGTA 0.3, HEPES 20, MgATP 4, and Na_3_GTP 0.3. Osmolality was adjusted to 290-295 mOsm and pH was buffered at 7.25–7.35. Series resistance was monitored and not compensated, and cells in which series resistance varied by 25% during a recording session were discarded. Synaptic responses were collected with a Multiclamp 700B amplifier (Axon Instruments, Foster City, CA, United States), filtered at 2 kHz, and digitized at 10 kHz. All recordings were performed at RT. A total of 100–300 consecutive miniature events were semi-automatically detected by off-line analysis using customized software Igor Pro (Wavemetrics) using a threshold of 6 pA. All mIPSC events were visually inspected to ensure that they were mIPSCs during analysis, and non-mIPSC traces were discarded. All pharmacological reagents were purchased from Abcam, and other chemicals were purchased from Sigma.

### Immunocytochemistry

The immunocytochemistry procedure in neurons was performed as described previously.(15) The indicated plasmids (SLITRK3-WT-Myc, SLITRK3-E606X-Myc or SLITRK3-C566R-Myc together with pCAGGS-IRES-GFP (SLITRK3:GFP=9:1)) were transfected into cultured hippocampal neurons at DIV 14-15 using Lipofectamine 3000 (ThermoFisher, L3000015). After 48 hours, the transfected cells grown on coverslips were gently rinsed with 1x PBS and fixed with 4% paraformaldehyde (PFA) and 4% sucrose in 1x PBS solution for 15 min at RT, followed by permeabilization with 0.2% TritonX-100 in 1x PBS for 15 min. Neurons were subsequently blocked with 5% normal goat serum in 1x PBS for 1 h and then incubated with primary antibodies as follows: anti-GFP (Cell signaling, #2555S, R), anti-Gephyrin (1:500, 147018, Synaptic Systems), and anti-vGAT (1:500, 131004, Synaptic Systems) in 1x PBS solutions overnight at 4°C. Cells were washed three times with 1x PBS and then incubated with Alexa Fluor 488, 555, or 647-conjugated IgG for 1 hour. Coverslips were washed three times with 1x PBS and mounted with Fluoromount-G (Southern Biotech) for imaging acquisition.

### Image Acquisition and Analysis

Fluorescence images were acquired with a Zeiss LSM 880 laser scanning confocal microscope using a 63x oil-immersion objective lens (numerical aperture 1.4). For gephyrin puncta density analysis, confocal images from 1 to 3 secondary or tertiary dendrites (35 μm in length) per neuron from at least ten neurons in each group were collected and quantified by counting the number of puncta per 10 μm dendrites with ImageJ puncta analyzer program. Thresholds were set at 3 SDs above the mean staining intensity of six nearby regions in the same visual field. Thresholded images present a fixed intensity for all pixels above the threshold after having removed all of those below. Labeled puncta were defined as areas containing at least four contiguous pixels after thresholding.

### Animal experiments

All animal experiments were approved by Animal Experiment Committees at the RIKEN Brain Science Institute and Animal Care and Use Committee of Nagasaki University. Further, they were conducted following the guidelines for animal experimentation in RIKEN and Nagasaki University. Same sex litter mates were housed together with two to four mice per cage. All mice were maintained on a regular diurnal lighting cycle (12:12 light:dark) with ad libitum access to food (CE-2, CLEA Japan) and water. Cellulose bedding material (SAFE comfort natura, Oriental Yeast, Japan) was used as bedding. The sample sizes for each experiment were determined such that the power and significance in the two-sided test were 80 and 5%, respectively. However, the number of samples from the animals was minimized empirically. Control groups consisted of littermates of the same sex. One or two litters were reared in a cage. Total number of mice and cages allocated to each experiment are summarized in Supplementary Table 1. Behavioral, MRI, and interneuron analysis were carried out using B6N1/N2 mice. No inclusion/exclusion criteria have been set before the study. All data were collected according to the randomly defined order of genotyping (tail cut). Potential confounders were not controlled. All animal experiments were carried out by experimenters who were blinded to animal identity. We have not set humane endpoints for maintaining the animal colony because some mice died suddenly.

### Behavioral analysis

Adult male Slitrk3 KO (*Slitrk3^tm1Jaru^/Slitrk3^tm1Jaru^*, *Slitrk3^-/-^*) and WT mice (*Slitrk3^+/+^*, 8–32-week-old, littermates from mated heterozygotes) were used for behavioral tests. Male mice were used to avoid effects of estrous cycles on behavioral phenotypes in females (25). Mice were housed in a 12:12 h light-dark cycle, with the dark cycle occurring from 20:00 to 8:00, and behavioral experiments were carried out between 10:00 and 17:00. Please see the Supplemental data section for a description of these behavioral experiments.

### Magnetic resonance imaging (MRI) based volumetric analysis

MRI images of the adult male mice were acquired by subjecting anesthetized mice to an MRI scan using a vertical bore 9.4-T Bruker AVANCE 400WB imaging spectrometer (Bruker BioSpin, Rheinstetten, Germany). Animals were anesthetized with 3% and 1.5% isoflurane in air (2 L/min flow rate) for induction and maintenance, respectively. MRI images were obtained by using the FISP-3D protocol of Paravision software 5.0, by setting the following parameter values: Effective TE, 4.0 ms; TR, 8.0 ms; Flip angle, 15 degree; Average number, 5; Acquisition Matrix, 256 × 256 × 256, FOV, 25.6 × 25.6 × 25.6 mm. Regional volumetric changes were measured by tensor-based morphometry (26). The images were subjected to non-uniformity correction and intensity standardization, before a multi-iteration group-wise registration, including 5 iterations of affine registration and 3 iterations of non-rigid registration, was performed. The Jacobian determinant for each voxel, which indicates its relative expansion or contraction as a result of the transformation, was derived from the deformation fields from the final round of non-rigid registration. This was followed by voxelwise two-tailed t-tests using a general linear model to compare the KO and WT groups, with total brain volume as a covariate. Non-uniformity correction (27) and image registration were performed using the NifTK software(28). Intensity standardization (29) and statistical analysis were performed using custom MatLab scripts.

### Interneuron counting

Mice were anesthetized with inhalation of isoflurane. Cardiac perfusion was performed with 4% paraformaldehyde and 0.1 M sodium phosphate (pH 7.4). Excised brains were fixed in the same fixative for 4 h at room temperature with gentle agitation. Tissue blocks were cryoprotected with 20% sucrose in PBS at 4°C overnight and frozen in OCT compound (Sakura Finetek). Cryosection was performed using a CM3050 cryostat (Leica Biosystems) at a thickness of 12 μm. After sectioning, sections were immersed in PBS(-) for 5 min, followed by 0.3% hydrogen peroxide in methanol for 10 min at room temperature. After immersion in PBS(-), the sections were blocked with 1% skim milk (Difco), 2% normal goat or donkey serum and 0.1% Triton X-100 in PBS (-) at room temperature for 1 h and reacted with mouse anti-parvalbumin monoclonal antibody (Sigma-Aldrich, P3088, 1:4000), goat anti-somatostatin antibody (Santa-Cruz, SC7819, 1:250), or sheep anti-neuropeptide Y antibody (Millipore, AB1583, 1:500) at 4°C for 0.5–2.5 days. The bound antibody was detected by VECTASTAIN Elite ABC kits (Vector Laboratories) and 3,3’-diaminobenzidine (DOJINDO). For the immunostaining using anti-somatostatin antibody or anti-neuropeptide Y antibody, the sections were autoclaved (105°C for 5 min) in 10 mM sodium citrate (pH 6.0) before the peroxidase-methanol treatment for antigen retrieval. Images were acquired with NDP slide scanner (Hamamatsu Photonics). For quantitative analyses, all stained images were taken with the same settings, and manually counted by observers who were blinded to the genotypes.

### Protein-protein network analysis

The interacting proteins with the SLITRK3 was constructed using the STRING server (https://string-db.org/) to identify both known and predicted protein-protein interactions. The network parameters (clusters, hubs, cliques and communities) were utilized to analyze the protein-protein interaction network. The clusters, and groups of interacting proteins (nodes) in the network were calculated. SLITRK3 was used as the seed in the construction of a protein-protein network to identify proteins that might involve in the interaction visualized by STRING and Cytoscape v.3.9.1 (30). The SLITRK3 node is the query and remaining nodes are relevant proteins having most connectivity with the query protein in the network of 11 proteins. In the bilateral network, the ‘nodes’ represent the protein targets and ‘edges’ represent the interactions of proteins.

PPI collection using PINOT, filtering and visualisation of networks: The direct interactors of SLITRK3 were collected through PINOT (31) on 1/11/2022 (Settings: Organism: Homo sapiens; Filter level: Stringent). As the data were limited all interactions were retained, and complemented by PPIs identified in the literature through manual curation (by querying SLITRK3 in PubMed on 26/8/2022). The manually curated interactions were of two categories: human data, or mouse data. The first were considered of higher confidence. The second layer interactome of SLITRK3 was built based on the results from PINOT and the high confidence interactors from the manual curation, using PINOT on 1/11/2022 (Settings: Organism: Homo sapiens; Filter level: Stringent). From the indirect interactors of SLITRK3 only those with FS higher than 2 were retained in the network. The PPINs were visualised using Cytoscape (v3.9.0)(30).

Enrichment and grouping: The 2^nd^ layer SLITRK3 network was analysed using enrichment through g:Profiler (32) on 2/11/2022 (version e107_eg54_p17_bf42210) (Settings: Statistical domain scope: Only annotated genes; Significance threshold: Bonferroni correction, User threshold: 0.05, No electronic GO annotations). The significant GOBP terms were grouped in semantic classes based on semantic similarity using in house dictionaries of the Manzoni lab (31). The semantic classes were grouped further in functional groups. The functional groups named General and Metabolism were not included in the analysis, as done previously (33). Neuronal-related terms were observed in multiple functional groups, so text mining using the keys “nerv”, “neur”, “brai”, and “synap”, was used to identify these terms in our results. The calculation of the enrichment ratios of the neuron-related words and the statistical analysis were calculated, as previously described (33). The p value was calculated using the Excel function: =2*(1-NORM.DIST(x, mean,sd,TRUE)).

### Protein Modeling of WT and mutant SLITRK3

There are currently no 3-D crystal structures of the SLITRK3 protein available, even no identified templates were found for comparative modeling. Therefore, I-TASSER Structure Prediction server was used for predicting the protein tertiary structure (34) as described in Supplementary section.

#### Data availability

The data that support the findings of this study are available from the corresponding author.

## Results

### Clinical findings

Family 1 consists of two 7-year-old (F1:II-1) and 6-year-old (F1:II-2) boys, and a 5-year-old girl (F1:II-3) from a consanguineous family of Pakistani descent (parents are first cousins) (Fig.1). Family history was unremarkable, except for one prior spontaneous miscarriage. The pedigree suggested an autosomal recessive inheritance. All presented with generalized tonic-clonic seizures since a young age (6 months for F1:II-1, 24 months for F1:II-2, and 18 months for F1:II-3) and extensor spasms in F1:II-2. No significant evolution of the epileptic phenotype emerged over time and seizures were controlled by antiepileptic drugs alone or in combination. All the patients developed progressive neurodevelopmental delay, with poor or absent speech and severe intellectual disability. F1:II-2 and F1:II-3 have short stature; 94 cm (<0.4^th^ centile) 84 cm (<0.4^th^ centile), respectively. F1:II-3 also has microcephaly at 44 cm (<0.4^th^ centile). EEG showed left parietal focal epileptogenic activity in F1:II-1 and diffuse encephalopathy in his two siblings. Ophthalmological examination in F1:II-1 showed retinal pigment epithelium abnormalities. Upon neurological examination, all three siblings presented with hyperactivity, whereas spasticity, brisk DTR and moderate weakness were observed in F1:II-1 and F1:II-2. Brain MRI in F1:II-1 was unremarkable.

Family 2 consists of a 5-year-old boy of (F2:II-1) of American origin (Fig.1). Pregnancy was complicated by intrauterine growth restriction (IUGR) but the child was born full term. He first presented with febrile seizures at 3 years of age accompanied by generalized fever (102°F, 38.9 C) with stiffness, focal shaking and limping, and which lasted less than one minute. Six months later he experienced tonic-clonic seizures with post-ictal symptoms. Since then, he experienced both febrile and non-febrile seizures, which manifested as generalized tonic-clonic episodes. Overnight video EEG at the age of 2 years showed background slowing, occasionally max left temporal, multifocal interictal epileptiform discharges (IEDs), max right central, often admixed with sleep architecture (Supplementary Fig. 1). Seizures were controlled by Levetiracetam alone. Brain MRI at the age of 2 years showed multiple subcortical white matter T2-hyperintensities without enhancement or mass effect. These lesions were non-specific and the differential diagnosis was long including gliosis, demyelination, dysmyelination, Lyme, vasculitis and chronic ischemia. There was optic nerve head flattening suggestive of papilledema. There were no developmental concerns, and a motor skill assessment revealed no atrophy or fasciculation, but a mild hypotonia was observed. Ophthalmology evaluated him and dilated exam showed normal healthy optic nerves without any edema. The patient was alert and interactive, he made good eye contact, and followed commands. Speech was fluent and language was appropriate for his age. Recent and remote memory are grossly intact.

Family 3 consists of a 16-year-old girl (F3:II-1) of African American origin (Fig. 1). The pregnancy was complicated by IUGR but she was born full term. Parents were concerned about motor delays very early on but became even more concerned around the age of 9-12 months. She was diagnosed with epilepsy, severe global developmental delay and intellectual disability. She rolled over at 6 months but did not meet any other motor milestones until after 2 years old and she did not walk until 6 years old. She currently remains nonverbal and does not use signs; she communicates via vocalizations and walking to what she wants. She started having staring spells at around 9-12 months old. These progressed to generalized tonic-clonic seizures lasting up to 2 minutes with post-ictal symptoms. Seizures were controlled with Leviteracetam and Clonazepam. Her last seizure occurred at the age of 14.5 years. F3:II-1 has short stature (<0.4^th^ centile), microcephaly at 47 cm (<0.4^th^ centile), and dysmorphic features (larger ears, prominent nose, wide mouth, short philtrum, everted upper lip, camptodactyly, flat feet). She has also been diagnosed with cortical vision impairment and nystagmus. The mother reported significant behavioural issues including breath holding, difficulty staying asleep, teeth grinding, hand rubbing, laughing fits, and crying spells. Brain MRI at the age of 11 months showed mild ventricular enlargement and prominent sulci suggestive of decreased brain volume, thin corpus callosum, and an irregular border to the ventricular system, raising the possibility of periventricular leukomalacia. Other clinical features included hypotonia and spasticity. Details clinical details of the patients identified in our cohort can be seen in the Appendix (Supplementary Table).

### Genetic findings

In total, 83,572,847 (II-1) and 81,527,162 (II-2) unique reads were generated by WES for F1:II-1) and 6-year-old (F1:II-2). After applying the previously mentioned filtering criteria, no plausible shared compound heterozygous variants were identified by WES; there was however a gene carrying a novel (likely) pathogenic variant, according to ACMG variant interpretation guidelines (21), which was homozygous in all three probands in family 1 (Fig. 1). A homozygous frameshift deletion in *SLITRK3* (NM_014926: c.1816G>T, p.Glu606*) emerged as the most likely explanation for the disease pathogenesis; this is also supported by a more severe impact of the mutation on protein function (truncating vs. missense) and an existing functional report previously linking cell adhesion molecules (CAM) such as SLITRK3 to synaptogenesis and GABAergic synapse development (13). The variant was residing within the largest region of homozygosity (out of two) in all three affected individuals of family A (Figure 1C). Segregation analysis performed by traditional Sanger sequencing confirmed the homozygous variant in the three affected siblings and heterozygous in both their parents. The identified *SLITRK3* homozygous variant was submitted to the Leiden Open Variation Database (www.lovd.nl/; variant ID #0000693802). Through clinical whole exome sequencing carried out at GeneDx, a *de-novo SLITRK3* variant (c.1696T>C, p.Cys566Arg) was identified for family 2 and a *de-novo* mosaic frameshift variant (c.660_684del25, p.Gly221Trp*fs*\*15) in family 3.

We further searched data from the 100,000 Genomes Project (Genomics England) (35) for families where affected individuals harboured monoallelic or biallelic variants in *SLITRK3*. All genomes from probands and affected family members (n=15,790) recruited under the Neurology Disease group and Mitochondrial disorders in the 100KGP were annotated and analysed for *SLITRK3* variants. Within this cohort, 2,638 individuals were recruited under the “Inherited Epilepsy Syndromes”. After filtering the dataset for *de-novo* variants, seven possible cases were identified. Weak phenotypic overlap and similarity across the seven hits led to the exclusion of these individuals from this cohort.

### Molecular findings

We performed the Western blot assay using an anti-SLITRK3 antibody recognizing SLITRK3 C-terminus and an anti-Myc antibody to identify the expression pattern of SLITRK3-WT-Myc, SLITRK3-C566R-Myc and SLITRK3-E606X-Myc in HEK293T cells (Fig. 2A). We found that the molecular weight of SLITRK3-C566R-Myc was slightly higher than WT, likely due to the altered glycosylation of the mutant. As the truncated form of the SLITRK3 mutant, For E606X, which is a nonsense mutation, leading to a loss of part of the transmembrane domain, and the entire C-terminus of SLITRK3. Indeed, using the SLITRK3 antibody, the E606X mutation could not be detected (Fig. 2A).Thus, we probed the E606X mutant using an anti-Myc antibody and found that the SLITRK3-E606X mutation resulted in a truncating molecule with lower molecular weight than the SLITRK3-WT protein.

**Fig 2.**
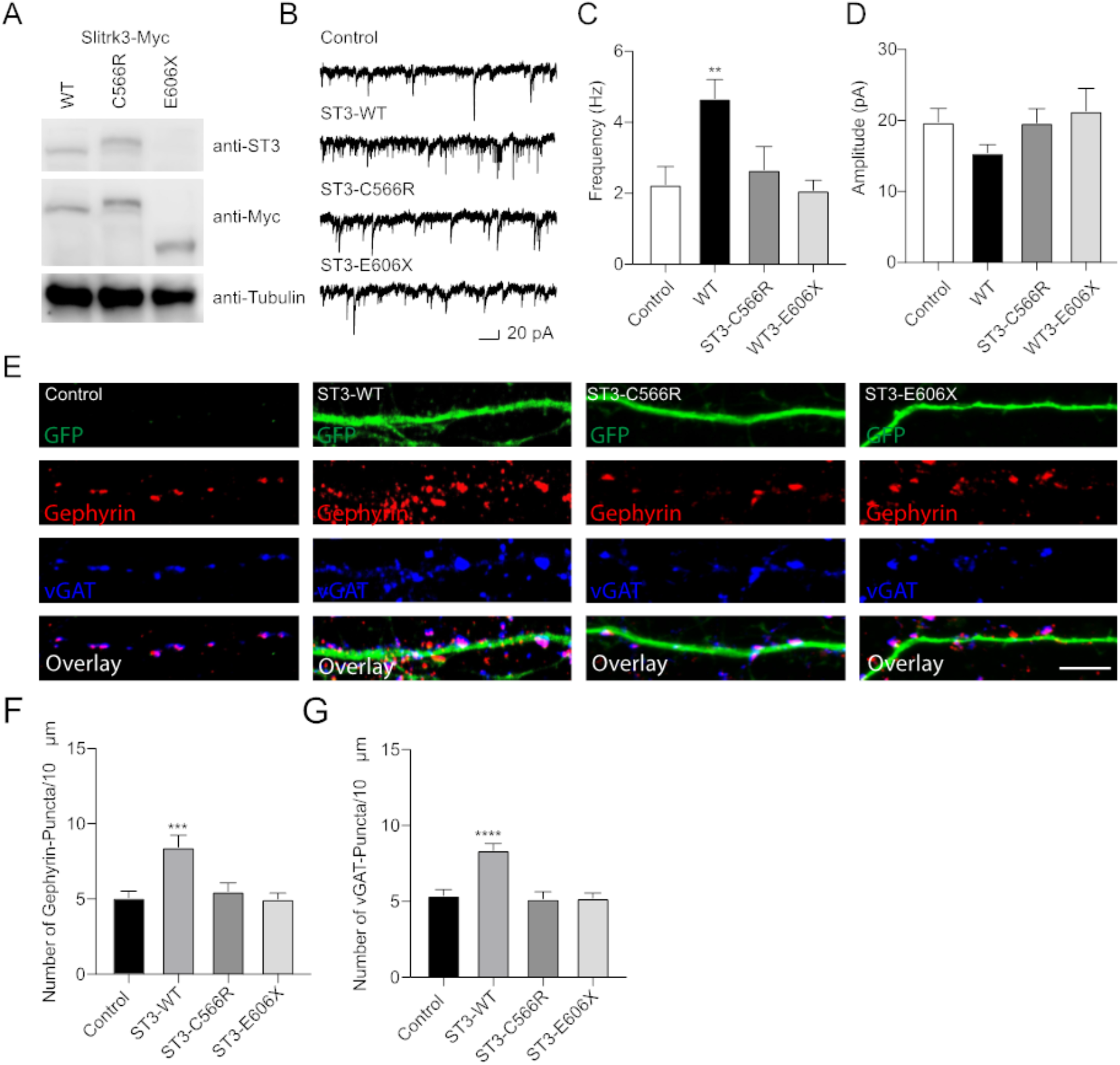
Functional effect of SLITRK3 variants on inhibitory synapses. (A) Identification of SLITRK3-WT, SLITRK3-C566R and SLITRK3-E606X expression in HEK-293 cells. Cell lysates from HEK-293 cells transfected with SLITRK3-WT-Myc, SLITRK3-C566R-Myc or SLITRK3-E606X-Myc were blotted with anti-SLITRK3, anti-Myc and anti-tubulin antibodies. N = 3 independent repeats. (B-D) Human mutants in SLITRK3 is essential for GABAergic synaptic transmission. (A) mIPSC recording showed that overexpression of human WT-SLITRK3 significantly increased mIPSC frequency, whereas overexpression of the SLITRK3-C566R mutant had no effect on the frequency of mIPSCs in cultured hippocampal neurons. Insets showed the mean ± SEM of mIPSC frequency and amplitude [Frequency (Hz): Control, 2.25 ± 0.50, n = 10; WT-SLITRK3, 4.67 ± 0.54, n = 9; SLITRK3-C566R, 2.67 ± 0.65, n=11; SLITRK3-E606X, 2.08 ± 0.28, n=10. One-way ANOVA test, ** p = 0.0058; Mann-Whitney test, Control vs SLITRK3-WT: ** p = 0.004; Control vs SLITRK3-C566R: p = 0.918; Control vs SLITRK3-E606X: p = 0.579. Amplitude (pA): Control, 19.770 ± 1.91, n = 10; SLITRK3-WT, 15.44 ± 1.14, n = 9; SLITRK3-C566R, 19.65 ± 2.02, n=11; SLITRK3-E606X, 21.30 ± 3.22, n=10. One-way ANOVA test, p = 0.33; Mann-Whitney test, Control vs SLITRK3-WT: p = 0.08; Control vs SLITRK3-C566R: p = 0.76; Control vs SLITRK3-E606X: p = 0.91. N = 5-6] Scale bar, 20 pA and 1 s. N = 6 independent repeats. (E-G) Identification of C566R and E606X in SLITRK3 that is important for GABAergic synapse density in hippocampal neurons. Representative images of dendrites and quantification analysis showed that overexpression of human SLITRK3-WT significantly increased vGAT and gephyrin puncta density in cultured hippocampal neurons, whereas overexpression of human SLITRK3-C566R and SLITRK3-E606X, did not change vGAT or gephyrin puncta density [vGAT: Control: 5.38 ± 0.39, n = 19, SLITRK3-WT: 8.36 ± 0.44, n = 17, SLITRK3-C566R: 5.17 ± 0.46, n = 16, SLITRK3-E606X: 5.22 ± 0.33. One-way ANOVA test, ****p < 0.0001, F = 12.22; Mann-Whitney test, Control vs SLITRK3-WT: **** p < 0.0001; Control vs SLITRK3-C566R: p = 0.62; Control vs SLITRK3-E606X: p = 0.66. Gephyrin: Control: 5.07 ± 0.45, n = 19, SLITRK3-WT: 8.44 ± 0.80, n = 17, SLITRK3-C566R: 5.50 ± 0.58, n = 16, SLITRK3-E606X: 5.00 ± 0.40. One-way ANOVA test, ***p = 0.0001, F = 8.038; Mann-Whitney test, Control vs SLITRK3-WT: ** p = 0.002; Control vs SLITRK3-C566R: p = 0.68; Control vs SLITRK3-E606X: p = 0.90]. N = 4 independent repeats. (SLITRK3 denoted as ST3 on figure).

To study the functional effect of these mutations on inhibitory synapses, we conducted the whole-cell recordings to measure miniature inhibitory postsynaptic currents (mIPSCs) in cultured hippocampal neurons (DIV14) overexpressing human WT-SLITRK3-Myc, SLITRK3-C566R-Myc or SLITRK3-E606X-Myc mutants (also simultaneously expressing GFP). In neurons overexpressing WT-SLITRK3, the frequency, but not amplitude, of mIPSCs was significantly increased (Fig. 2B–D), similar to an early study (15). In contrast, in neurons overexpressing SLITRK3-C566R, GABAergic transmission had no change compared with the control cells showing that the missense mutation of C566R essentially abolished the effect of SLITRK3 overexpression on GABAergic transmission (Fig. 2B–D). Thus, SLITRK3-C566 plays a critical role in the regulation of inhibitory transmission. In addition, the mIPSC frequency and amplitude had no change in neurons expressing SLITRK3-E606X mutant compared with the control cells (Fig. 2B–D), suggesting that premature termination of SLITRK3 at E606 inactivates the function of SLITRK3 in promoting GABAergic transmission.

To further examine the role of human SLITRK3 mutants, we investigated the function of these mutants in the regulation of GABAergic synapses in hippocampal neuronal cultures, as SLITRK3 is a key inhibitory synaptic cell adhesion molecule. To this end, we over-expressed SLITRK3-WT-Myc, SLITRK3-C566R-Myc or SLITRK3-E606X-Myc together with pCAGGs-IRES-GFP in dissociated hippocampal neurons, and examined the densities of vGAT and gephyrin, the inhibitory pre- and post-synaptic markers, respectively, in neuronal dendrites. The GFP expression pattern helped delineate the transfected neuronal dendrites. We found that both vGAT and gephyrin in neuronal dendrites were significantly increased in hippocampal neurons overexpressing SLITRK3-WT-Myc (Fig. 2E–G), consistent with the recent study (15). In contrast, overexpression of human SLITRK3 mutants of SLITRK3-C566R-Myc or SLITRK3-E606X-Myc did not change the densities of vGAT and gephyrin in neuronal dendrites (Fig. 2E–G), showing that these two mutations abolish the ability of SLITRK3 in enhancing GABAergic synapse development.

### Animal studies

In a previous study, SLITRK3 KO mice were shown to have reduced inhibitory synapse density and to exhibit increased seizure susceptibility and abnormal epileptiform activity in electroencephalogram (12). However, we have not observed either spontaneous or handling-induced seizure-like behavioral abnormalities during their maintenance or behavioral tests. To better understand the biological role of SLITRK3, we analyzed the developmental behaviour and phenotypes of SLITRK3 KO (*SLITRK3^-/-^*) mice. In the postnatal development, SLITRK3 KO mice exhibited incomplete lethality. Genotyping at adolescent period indicated that 59% (B6N1 /N2, KO mice generated in 129P2 strain background backcrossed to C57BL/6 strain once or twice) or 73% (B6N8, similarly backcrossed eight times) of the SLITRK3 KO mice died before 4 weeks-old (Table 1). There was a significant difference in the genotype ratios between the B6N1/N2 group and B6N8 group (P = 0.0095 in **χ**^2^-test, Table 1), suggesting that the lethality was affected by the difference in genetic backgrounds. In the longitudinal analysis of B6N8 mice, 7 out of 9 SLITRK3 KO mice died on P20 or P21, and body weight was significantly lower than SLITRK3 WT both at 2 weeks-old (−29%, P = 0.0022) and 3 weeks-old stages (−45%, P = 0.00048) (Fig. 3). On the other hand, B6N1/N2 group mice did not show significant body weight gain even at adult stages (Fig. 3). Adult SLITRK3 KO (B6N1/N2) mice are known to exhibit enhanced susceptibility to pentylenetetrazole-induced seizure and the appearance of spontaneous epileptiform EEG (12). However, we rarely observed either spontaneous or handling-induced seizure-like behavioral abnormalities during their maintenance or behavioral tests, and little is known about the other neurological abnormalities. We then investigated their behavioral abnormalities. As a result, SLITRK3 KO mice showed higher motor activities than WT littermates in several tests. In their homecages, higher spontaneous activities were observed both at light and dark phases (Fig. 4A). On rotating rods, SLITRK3 KO mice stayed longer time at first round of the tests (Fig. 4B). On elevated plus maze apparatus, SLITRK3 KO mice moved longer distances (Fig. 4C). In forced swimming, SLITRK3 KO mice showed less immobility (Fig. 4D). The forced swimming test is often used as to assess the effect of antidepressant. However, other behavioral tests used to assess mood-associated behaviors such as immobile time in tail suspension test or marble burying test did not show differences between WT and KO (Supplementary Fig. 2).

**Fig 3.**
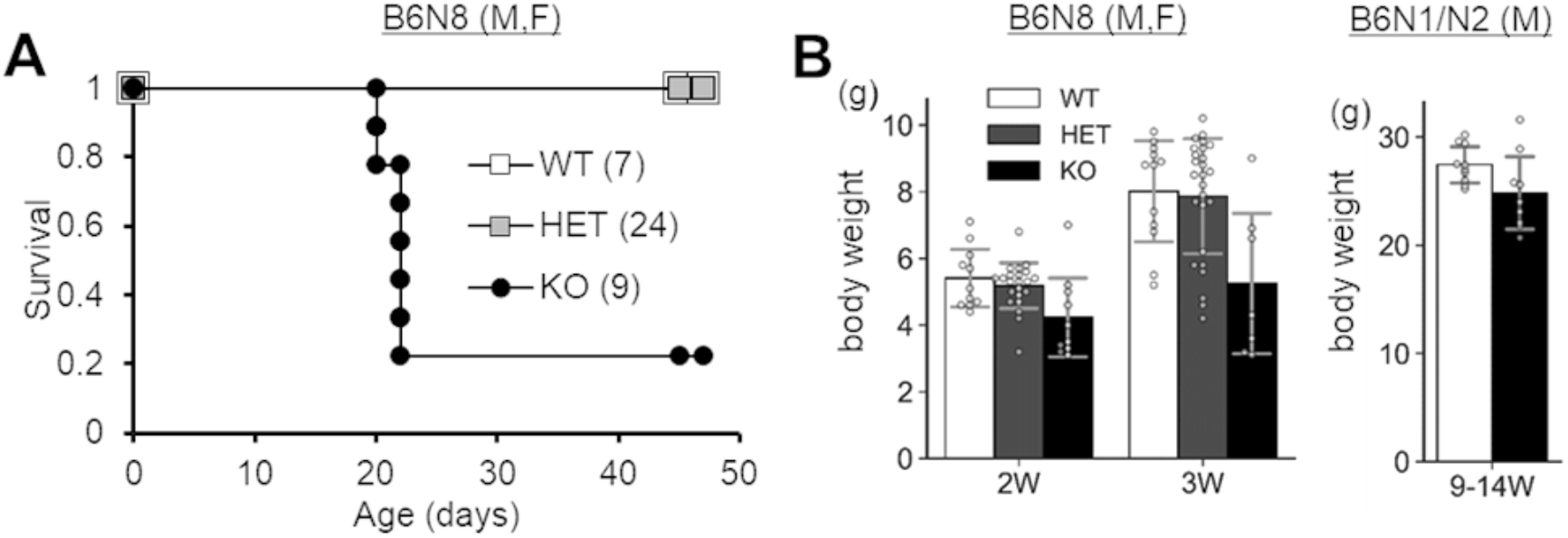
Survival and development of SLITRK3 KO mice. Genotype counts and percentages of progenies generated by inter-heterozygous mating. Genotyping was done for B6N1/N2 progenies and B6N8 progenies at their ages of 4.1 to 7.0 weeks-old.

**Fig 4.**
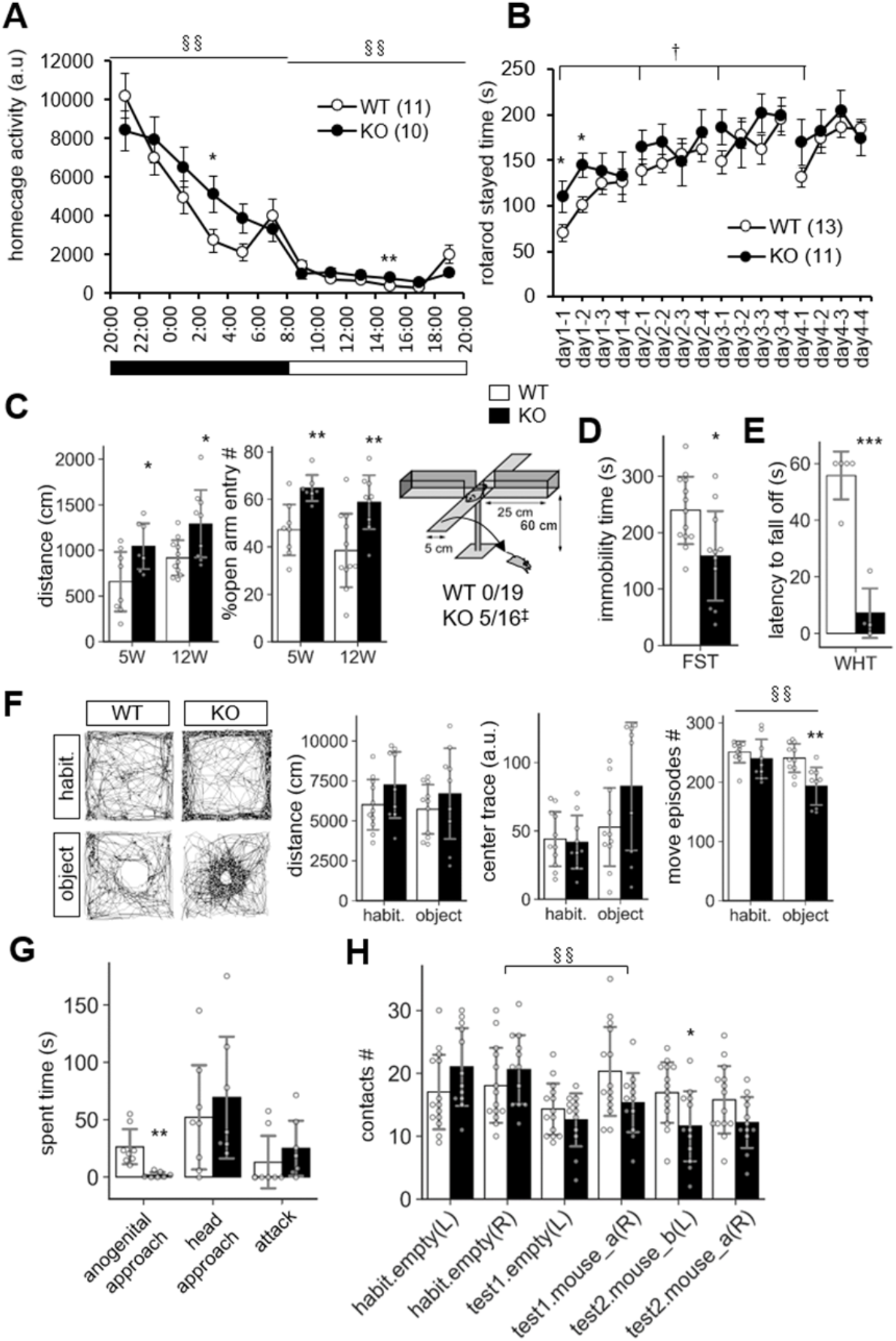
Behavioral abnormalities of SLITRK3 KO mice. (A) Spontaneous activities in homecages. §§, *P_(genotype × time)_* < 0.01; in two-way repeated-measures ANOVA (time and genotype as main factors). (B) Rotarod test. The tests are carried out daily on day 1 to day 4 (4 trials per a day). The stayed time on accelerating rotarod is measured. †, *P_(genotype)_* < 0.05; in twoway repeated-measures ANOVA (1st trial and genotype as main factors). (C) Elevated plus maze test. (*Left*) Total distance traveled. (*Middle*) Percentages of the numbers of open arm entries to total arm entries. (*Right*) Falling off from open arms. Values indicate (number of fallen mice)/(total number of mice). ‡, p < 0.05 in Fisher’s exact test. (D) Immobility time in forced swim test (FST). Immobile time in the test was 10 mins. (E) Wire hanging test (WHT). Latency to fall off from wire mesh after flipping was measured. (F) Novel object approach test. In habituation session (*habit*.), mice were left in an open field for 15 min. In novel object session (*object*), mice were left in an open field with an unfamiliar object (inverted paper cup) for 15 min. Traveled distance (*left*), trace (*middle*), and number of move episode (*right*) during each session were measured. §§, *P_(session × time)_* < 0.01; in two-way repeated-measures ANOVA (session and genotype as main factors). (G) Resident intruder test. Times spent in approaching to the anogenital region (*anogenital approach*) or head region (*head approach*) of intruder mice, and that in biting to intruder mice (*attack*) were measured. (H) Social discrimination test. Graph indicates the number of contacts to cylinder cages with or without a mouse. The test consisted of three successive 15 min sessions: habituation session (*habit*.), two empty cages at left and right corners; *test1* session, one empty cage at left and a new mouse-containing cage at right; *test2* session, a new mouse containing cage at left and a familiar mouse containing cage at right. §§, *P_(session × time)_* < 0.01; in two-way repeated-measures ANOVA (session and genotype as main factors). (A-H) *, p < 0.05; **, p < 0.01; ***, p < 0.001 in unpaired two-tailed Student’s t-test. Mean values are indicated in all graphs. *Open bars and circles*, WT; *closed bars and circles*, KO. *Error bars*, SD (bar graphs), SEM (line graphs). *Gray circles on bar graphs* indicate the individual value of each mouse. The *numbers in parentheses* in line graphs indicate n (the number of mice) in each experimental group.

**Table 1.**
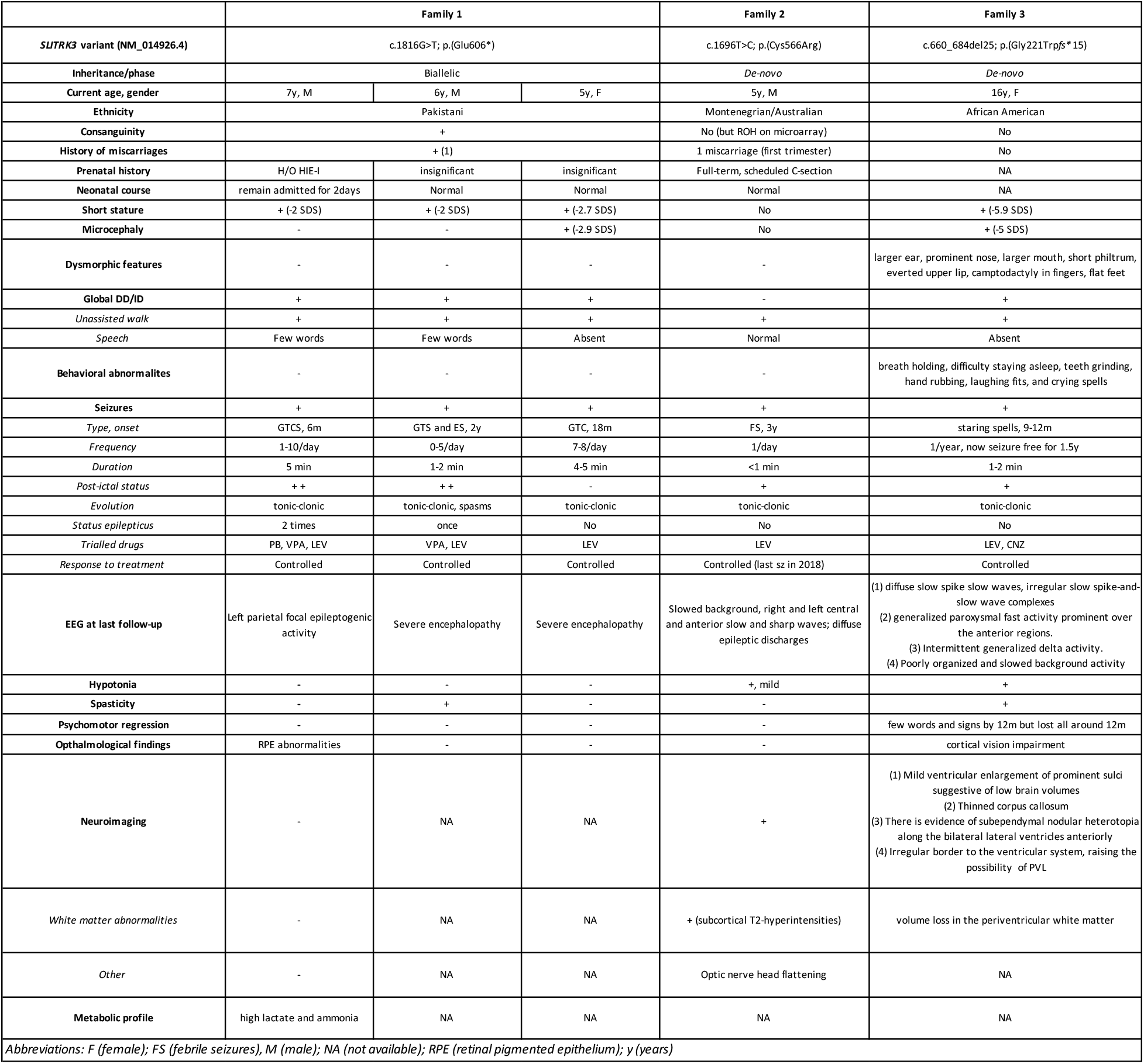
Summary of the clinical features of SLITRK3 patients.

Another characteristic feature included the increased open arm entries and falling from the elevated plus maze apparatus (Fig. 4C) and decreased stayed time in wire-hanging test (Fig. 4E). These signs were thought to reflect either reduced anxiety and/or impaired sensorimotor function to prevent falling off. There were no clear genotypes differences in anxiety-related behavioral indexes such as center-area stayed time in open field test and light-dark box transition test (Supplementary Fig. S2). On the other hand, we observed significant WT-KO genotype differences in some other sensorimotor function-related parameters such as increased freezing response after first conditioned-unconditioned stimulus in fear conditioning test and decreased auditory startle response to 90 dB and 95dB stimuli in auditory startle response test (Supplementary Fig. S2). Although the sensory modalities were different among the tests (i.e. vision-depth-height, nociception-foot-shock, auditory-noise), it seemed likely that SLITRK3 KO involves some sensory system dysfunction.

In addition, SLITRK3 KO mice showed different responses to both inanimate and animate objects. In case of inanimate object (a small cylinder) on open-field apparatus, SLITRK3 KO mice showed fewer move episode counts than WT mice (Fig. 4F). In reciprocal social interaction test (resident-intruder test), SLITRK3 KO mice (resident) showed fewer approach to the anogenital region of the unfamiliar (intruder) mice (Fig. 4G). In social discrimination test, SLITRK3 KO mice showed fewer approach to the cage with unfamiliar mice (Fig. 4H). Collectively, the behavioral abnormalities of SLITRK3 KO mice may be well categorized into the context-dependent higher motor activities, sensorimotor function abnormalities, and altered responses to unfamiliar objects. Otherwise, we can summarize these abnormalities in a broad term such as cognitive dysfunction.

Because a previous study showed defective inhibitory synapse formation on hippocampal pyramidal neurons in SLITRK3 KO mice, we investigated the interneurons numbers in hippocampus. It was found that numbers of parvalbumin-positive interneurons were reduced in hippocampal CA1, CA3, and dentate gyrus regions (Fig. 5A). On the other hand, numbers of somatostatin-positive neurons and neuropeptide Y-positive neurons were comparable between the two genotypes (Fig. 5A). We also quantified the parvalbumin-positive cell numbers in somatosensory cortex. However, there were no clear genotype-dependent differences (Fig. 5B). We also examined the gross morphology of SLITRK3 KO brains by MRI. The total brain volume was comparable between SLITRK3 WT and KO mice (Supplementary Fig. S3). There were no clear genotype-dependent regional volume changes in tensor-based morphometry analysis (Supplementary Fig. S3). These results indicated that SLITRK3 deficiency caused selective reduction in the parvalbumin-positive interneuron subset.

**Fig 5.**
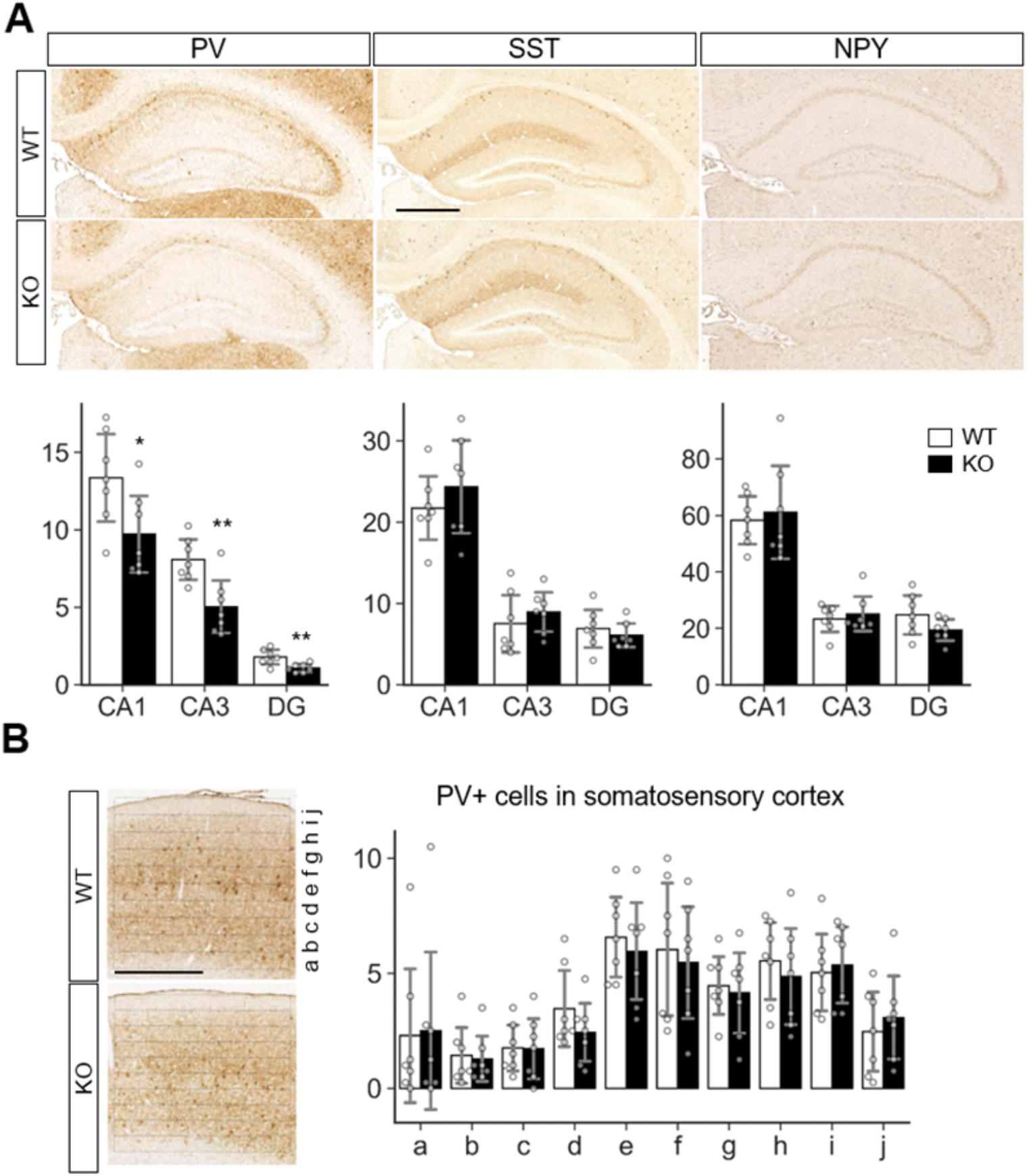
Quantitative analysis of inhibitory neuron subtypes in SLITRK3 KO brains. (A) Representative images for parvalbumin (PV), somatostatin (SST), and neuropeptide Y (NPY) immunostaining of the hippocampus CA1 stratum radiatum (CA1), CA3 stratum lucidum (CA3), dentate gyrus (DG) from P10-P16 WT (n = 7 mice) and KO (n = 7 mice) coronal sections. Sex-matched littermates WT and KO pairs (4 male pairs and 3 female pairs) were subjected to analysis. (B) Quantitative analysis of each inhibitory neuron subgroup. (C) PV positive neurons in cerebral cortex. PV-positive cells in WT or KO somatosensory cortices were counted in ten strips (a-j) parallel to the pia surface. (B, C) Immunopositive cells were counted by genotype-blinded observers. *Open bars*, WT; *closed bars*, KO. *Error bars*, SD. *Gray circles* indicate the individual values of each mouse, which are means of four independent images. *, p < 0.05; **, p < 0.01 in unpaired two-tailed Student’s t-test.

### In-silico modeling

Structural modelling of WT and mutant SLITRK3 proteins showed that in the WT protein, the Cys556 residue forms a disulfide bridge with the Cys591, which gets disrupted due to the heterozygous c.1696T>C, p.Cys566Arg variant. The mutated Arg566 residue forms an interaction with the vicinity residues involving the side chain. This results in disulfide bridge breaking and interaction of the Arg566 can alter the protein’s structural confirmation. Overall, there is similarity between WT and mutated structures but a significant position difference such as that of alpha helixes and β-sheets has been observed after running the energy minimization of these models. Nonsense variants such as the homozygous c.1816G>T, p.Glu606Ter is predicted to create a shorter protein, compared to normal SLITRK3, resulting in termination of the chain and loss of the cytoplasmic domain of the protein. Additional modeling of the frameshift mutation c.660_684del25, p.Gly221Trp*fs*\*15 with Chimera v.1.4. revealed the presence of only a chunk of the extracellular domain of the protein which is predicted to impede the protein’s normal function (Supplementary Fig. S4).

General protein-protein network analysis using STRING revealed different potential interactions with 11 other proteins such as other important CAMs i.e. NLGN2, NRXN2, PRPRS (Supplemenrary Fig. S5A). A more stringent protein-protein network analysis approach followed in which we included only human protein-protein interactions (PPIs) that were manually curated by experts, using the online tool PINOT (n=6 direct interactors). Manual curation of PPIs is producing data in which we can have higher confidence compared to other methods, e.g., text mining. However, it is more time consuming, potentially leading in the delay in the inclusion of recently published data. Since, SLITRK3 has limited literature, we decided to manually curate its literature and include additional PPIs in our network. This resulted in the identification of 4 additional interactions, of which only 1 was between two human proteins (i.e., NTRK3), while the rest were mouse proteins (i.e., Ptprd, Nlgn2, and Erbb4). The human interactors (n=7) were included in subsequent analysis, in which the 2^nd^ layer interactome was collected (i.e. interactions of the direct interactors of SLITRK3) (Supplementary Fig. S5B and C) and analysed further using enrichment of biological processes and pathways. The top resulted GOBP terms based on the adj p value as calculated by g:Profiler, were selected based on the distribution of the p values, as shown in Supplementary Fig. S5D (n=125). They had similar themes related to phosphorylation, intracellular signal transduction, ERBB signalling pathway, MAPK cascade, response to stress, cellular localization, cell migration, epidermal growth factor receptor signalling pathway, positive regulation of immune system process, cell death, vesicle-mediated transport, Fc receptor mediated stimulatory signalling pathway, receptor-mediated endocytosis, cell differentiation, cell adhesion, and apoptosis. Neuron-related terms were also observed and their enrichment ratios and p values of enrichment were calculated for “*nerv*”, “*neur*”, and “*synap*” (2.90, 0.018; 1.40, 0.037; and 1.29, 5.6×10^-9^), providing an additional indication of the role of SLITR3 and its interactome in the function of neurons. Pathway enrichment analysis highlighted pathways that could be implicated with SLITRK3’s physiological function, such as signal transduction, cytokine signalling in immune system, axon guidance, nervous system development and clathrin-mediated endocytosis (Supplementary Tables 2&3).

## Discussion

We identified de-novo heterozygous and bi-allelic mutations in *SLITRK3* in 5 individuals from 3 families, with a developmental epileptic encephalopathy phenotype. The clinical picture was predominated by early onset global developmental delay, seizures, intellectual disability and attention deficit–hyperactivity disorder, all of which pointed to a brain disorder. Early studies have shown that SLITRK3 is highly enriched at GABAergic inhibitory synapses and plays a critical role in the regulation of GABAergic synapse development (4, 12). Recently, mechanistic investigations have demonstrated that SLITRK3 is involved in GABAergic synaptogenesis at the late stage of development (13) and functions in adenosine receptor-mediated inhibitory synapse stabilization during development, a process depending on SLITRK3 Y969 (14, 15). Our data have now revealed that epilepsy-associated human mutations in SLITRK3 disrupt its function in regulating GABAergic synapses, leading to a number of neurological consequences in human patients.

In keeping with its essential role in GABAergic synapse development, SLITRK3 is widely expressed in all brain regions from early development, with highest levels found in the cerebral cortex and cerebellum (BRAINEAC database). The gene is intolerant to loss of function mutations in humans, with a pLi score of 0.94. Indeed, the mIPSC frequency and amplitude in hippocampal neurons expressing SLITRK3-E606X mutant had no changes compared with the control cells expressing WT SLITRK3 that increased inhibitory transmission. Consistently, expression of SLITRK3-E606X did not change dendritic densities of vGAT and gephyrin, while overexpression of WT SLITRK3 significantly increased GABAergic synapse development. These data indicate that E606X mutation abolishes the function of SLITRK3 in promoting GABAergic synapse development. We thus conclude that the phenotype in the index probands with the SLITRK3-E606X variant could be due to a loss of function of WT-SLITRK3. This could be explained by the fact that E606X is a truncation mutant lacking the important transmembrane domain and C-terminus.

Interestingly, expression of SLITRK3-C566R mutant also did not change mIPSCs and GABAergic density in hippocampal neurons, suggesting that the N-terminal residue, C566, is essential for SLITRK3 function in the regulation of GABAergic synapse development and that mutation of Cytosine 566 to Arginine completely inactivates SLITRK3-dependent promotion of GABAergic synapse development. In line with SLITRK3 being tolerant to missense mutations in humans, with a Z score of 0.59, we speculate that probably heterozygous de-novo non-synonymous variants likely affect the somewhere important?.Currently, the mechanisms underlying the role of C566 in regulating inhibitory synapses remain unclear. As SLITRK3 N-terminus is critical for trans-synaptic interaction with presynaptic PTPδ and cis interaction with postsynaptic NL2, it is possible that C566 might be important for SLITRK3 N-terminus mediated protein interactions crucial for its synaptogenic function.

Additionally, the overexpression studies showed no changes in dendritic densities of vGAT and gephyrin, suggesting these both variants SLITRK3-C566R and SLITRK3-E606X are essential to the function of SLITRK3 in the regulation of GABAergic synaptic densities. Together, we found the increased vGAT and gephyrin in neurons overexpressing SLITRK3-WT-Myc but no change of those in neurons overexpressing SLITRK3-C566R-Myc or SLITRK3-E606X-Myc, consistent with the change of mIPSC frequency in those transfected GFP-positive cultured hippocampal neurons.

In terms of seizure-associated phenotypes of SLITRK3 KO, we previously reported that enhanced susceptibility to pentylenetetrazole-induced seizure and the appearance of spontaneous epileptiform EEG (12). These results together with the developmental deficits shown in the current study strongly support the *SLITRK3* loss-of-function mutations as genetic causes of epileptic encephalopathy. In addition, the cognitive dysfunction in SLITRK3 KO mice might be associated with sensory or motor signs or intellectual disability presented in the patient with *SLITRK3* loss-of-function mutation.

PV interneurons in the hippocampus are mostly basket cells with fast-spiking actions at a high energy cost, and are highly vulnerable to stressors and have been implicated in many neuropsychiatric disorders (36). Impaired development or function of Parvalbumin-positive interneurons are associated with epilepsy in various animal models of epilepsy, as well as some genetic form of epilepsy in humans (37). In mice, selective silencing of hippocampal PV interneurons induces recurrent spontaneous limbic seizure (38). Therefore, reduced PV interneurons in SLITRK3 KO mice raise the possibility that deterioration of parvalbumin interneurons in the neurological signs of the patients with SLITRK3 loss-of-function mutations.

The clinical phenotype associated with the homozygous variant c.1816G>T, p.Glu606* (generalised tonic-clonic seizures, behavioural abnormalities, ADHD) correlates with absent protein levels and a shorter protein missing the cytoplasmic domain and could reflect impaired protein stability as suggested by *in-silico* structural modeling (Supplementary Fig S4). However, the clinical phenotype of the heterozygous variants c.1696T>C, p.Cys566Arg and c.660_684del25, p.Gly221Trp*fs*\*15 (periventricular leukomalacia with tonic-clonic seizures, motor delay, facial and behavioural abnormalities) point more towards altered structural confirmation of the proteins. Disulfide bonds can have profound effects on the folding pathway and the stability of a protein, thus increasing its suitability for existence in the extracellular milieu. However, due to the limited tendency of alpha-helical residues to form disulfide bridges in these two variants, we suspect that only a chunk of the extracellular domain is being formed, especially in the case of c.660_684del25, p.Gly221Trp*fs*\*15, which is predicted to be detrimental to normal protein function.

In conclusion, we have identified a neurodevelopmental disease with an early onset of symptoms that is variably associated with additional neurological features predominated by GDD/ID, ADHD, hypotonia, and epilepsy. Given that the synaptopathies are defined as brain disorders associated with synaptic dysfunction (39) and that the individuals presented in this study have clinical features overlapping those observed in individuals with synaptopathies (cognitive disorders such as intellectual disability, motor dysfunction such as ataxia and dystonia, epilepsy and psychiatric diseases such as ASD and ADHD), we coin the phenotypes associated with SLITRK3 variants as SLITRK3-related synaptopathy. Furthermore, variability in the effects of different SLITRK3 mutants under *in-vitro* conditions points toward mutation-specific mechanisms underlying the postsynaptic defect of the affected children, and this variability highlights a promising area of future research.

**Table 2.**
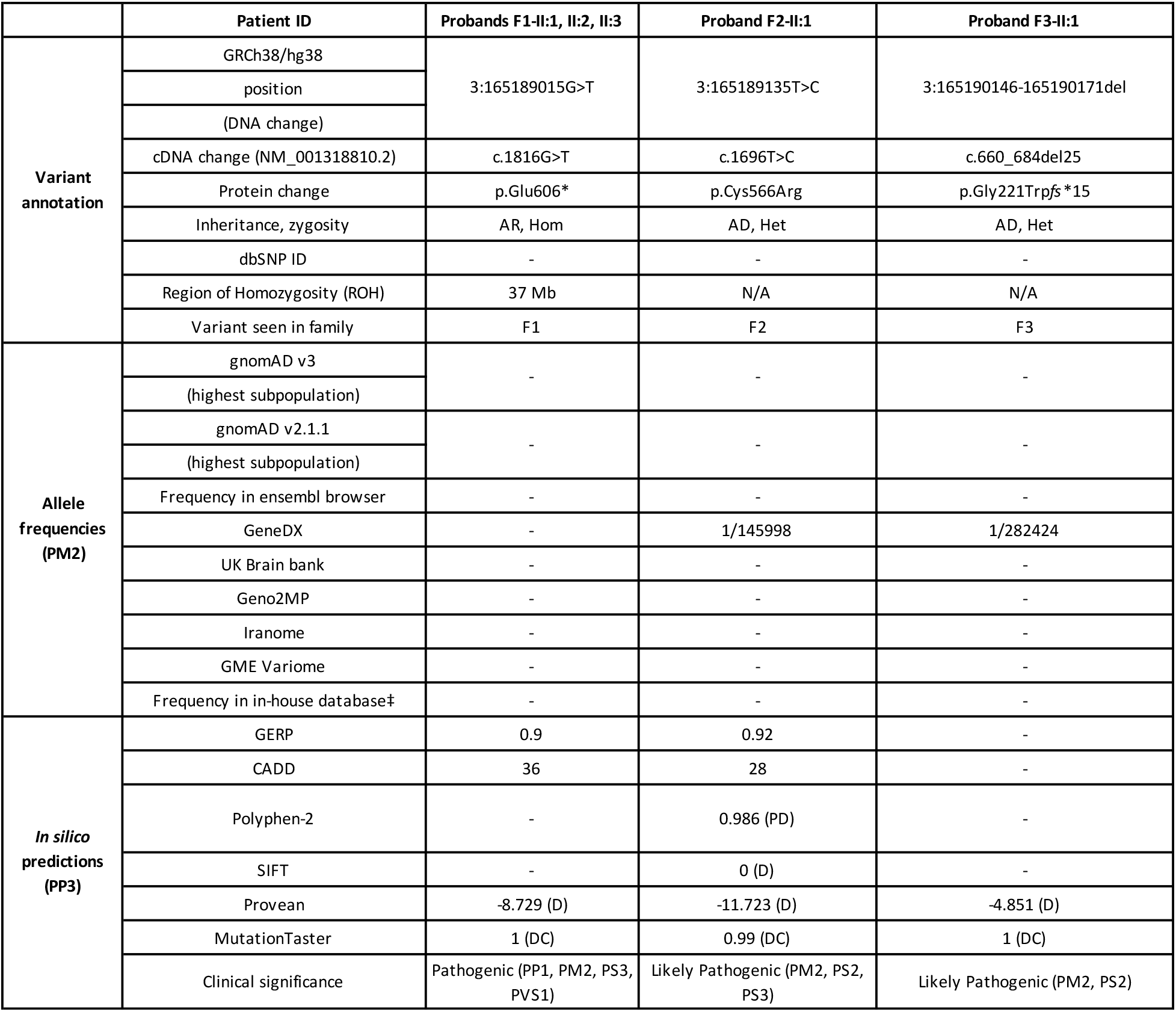
*SLITRK3* intragenic variants identified in our cohort.

**Table 3.** SLITRK3 KO mice exhibit postnatal lethality. Genotype counts and percentages of progenies generated by inter-heterozygous mating. Genotyping was done for B6N1/N2 progenies and B6N8 progenies at their ages of 4.1-7.0 weeks-old.

| genetic background | sex | ST3+/+ (WT) | ST3+/- (HET) | ST3-/- (KO) | total | χ<sup>2</sup>-test: Mendelian | χ<sup>2</sup>-test: B6N1-N2 |
| --- | --- | --- | --- | --- | --- | --- | --- |
| B6N1-N2 | Total | 280 (28.8%) | 527 (54.2%) | 166 (17.1%) | 973 (100%) | 5.43E-08 |  |
| B6N8 | Male | 30 (31.9%) | 54 (57.4%) | 10 (10.6%) | 94 (100%) | 5.00E-03 | 2.49E-01 |
|  | Female | 34 (42.5%) | 39 (48.8%) | 7 (8.8%) | 80 (100%) | 1.08E-04 | 1.16E-02 |
|  | Total | 64 (36.8%) | 93 (53.4%) | 17 (9.8%) | 174 (100%) | 2.03E-06 | 9.51E-03 |

## Acknowledgments

The original family was collected as part of the SYNaPS Study Group collaboration funded by The Wellcome Trust and strategic award (Synaptopathies) funding (WT093205 MA and WT104033AIA). This research was conducted as part of the Queen Square Genomics group at University College London, supported by the National Institute for Health Research University College London Hospitals Biomedical Research Centre.

## Author contributions

Stephanie Efthymiou: Conceptualization; Data curation; Formal analysis; Investigation;

Visualization; Methodology; Writing—original draft; Writing—review & editing

Wenyan Han: Data curation; Formal analysis; Investigation; Visualization; Methodology;

Writing—review & editing

Muhammad Ilyas: Data curation; Formal analysis;

Jun Li: Data curation; Formal analysis; Investigation; Visualization; Methodology;

Yichao Yu: Data curation; Formal analysis; Investigation; Methodology;

Marcello Scala: Methodology; Writing—original draft; Writing—review & editing

Nancy T. Malintan: Methodology; Writing—review & editing

Muhammad Ilyas: Data curation; Formal analysis;

Nikoleta Vavouraki: Data curation; Formal analysis;

Kshitij Mankad: Data curation; Formal analysis;

Reza Maroofian: Conceptualization; Methodology

Clarissa Rocca: Data curation; Formal analysis

Vincenzo Salpietro: Data curation; Investigation; Methodology

Shenela Lakhani: Data curation; Formal analysis; Investigation; Methodology

Eric J. Mallack: Data curation; Formal analysis; Investigation; Methodology

Timothy Blake Palculict: Data curation; Formal analysis;

Hong Li: Data curation; Formal analysis; Investigation; Methodology

Guojun Zhang: Data curation; Formal analysis; Investigation; Methodology

Faisal Zafar: Data curation; Formal analysis; Investigation; Methodology

Nuzhat Rana: Data curation; Formal analysis; Investigation; Methodology

Noriko Takashima: Data curation; Formal analysis; Investigation; Visualization; Methodology

Hayato Matsunaga: Data curation; Formal analysis; Investigation; Visualization; Methodology

Pasquale Striano: Data curation; Formal analysis; Investigation; Methodology

Mark F Lythgoe: Conceptualization

Jun Aruga: Conceptualization; Supervision; Funding acquisition; Writing—review & editing.

Wei Lu: Conceptualization; Supervision; Funding acquisition; Writing—review & editing.

Henry Houlden: Conceptualization; Supervision; Funding acquisition; Writing—review & editing.

