## Supplementary data for "Human mutations in *SLITRK3* implicated in GABAergic synapse development in mice"

**Supplemental data**

**Material and methods**

*Home cage activity measurement*

Spontaneous activity of mice in their home cage was measured using a 24-ch ABsystem 4.0 (Neuroscience, Tokyo, Japan). Cages were individually set into the compartments made of stainless steel in the negative breeding rack (JCL, Tokyo, Japan). A piezoelectric sensor was equipped on the ceiling of each compartment, and it detected movements of the mice. Home cage activity was measured for one week from the afternoon of the day of transferring to the behavioral laboratory (Day 1) until the next day of the week (Day 8).

*Rotarod test*

Rotarod evaluations were performed for both young (3–5-month-old) and aged (15 months) mice using a Rotarod Treadmill MK-610A Muromachi apparatus. Briefly, one habituation session was conducted 24 h before the beginning of the task, consisting of one practical trial of 2 min (4 rpm). The mice were then subjected to the paradigm for 4 consecutive days, each one consisting of four rotarod trials (with 1-min intervals between trials, allowing the animals to rest but avoiding them to become inactive). The rod accelerated from 4 to 40 rpm over 240 s and was maintained at 40 rpm for 60 s thereafter. The maximum trial duration was 300 s. Animals that fell off the rod or failed to turn one full revolution were returned to the cage.

*Elevated plus maze test*

Each mouse was individually put on the center platform facing to an open arm of the plus maze (closed arms: 25 x 5 x 15 cm (H); open arms 25 x 5 x 0.3 cm (H), arranged orthogonally 60 cm above the floor), and then allowed to move freely in the maze for 5 min. Total distance traveled, % time stayed in the open arms, % number of the open arm entry were measured as indices. Data was collected and analyzed using Image J EPM (O'Hara, Tokyo, Japan).

*Forced swimming test*

Each mouse was placed for 6 min in a glass cylinder (30 cm high, 10 cm in diameter) containing 20 cm of water maintained at 23–25˚C. The duration of immobility was recorded during the last 3 min of the test period.

*Open field and novel object tests*

Novel object approach test was performed in the open-field (OF) apparatus (50 × 50 × 40 [H] cm). A mouse was first placed in the OF with 70 lx illuminance for 15 min (habituation session). After the habituation session, the mouse was returned to its home cage and an inanimate object was placed in the center of the field. In the next test session, the mouse was placed again in the OF with the novel object. The large object was prepared by joining two paper cups by their openings (see Fig. 3A). Inside the bottom of one cup, a metal block was placed to give stability, and gray monotone and check-patterned printed papers were wrapped around the external surfaces of the cups. Each large object was discarded after use and a new object that had had no contact with the experimental animals was used. The mean time interval between two sessions was 4 min. The total distance traveled and % of time spent in the central area (30% of the field), which included the object and the area around it, were analyzed by using Image J OF4 (O'Hara). Contacts with the novel object were counted on the video records by an observer who was blind to the genotypes. Contact was defined as a forward movement toward the object and subsequent direct contact using the head

*Resident-intruder test*

Group-reared mice were kept in isolation for 5 days before the test. The test was carried out in a dark phase (0:30 to 2:30) in a chamber that keeps the cage under dim (infrared) light at 25 ˚C. Video recording from two opposite directions was initiated once the intruder mice had been gently placed in a vacant spot in the cage of the resident mice. The behaviors of the resident mouse were recorded for 10 min. The duration and number of times the resident mice spent sniffing, inactive contact with, and in pursuit and attack with the intruder mice were measured by observers who were blinded to the genotypes of the mice.

*Social discrimination test*

This test was performed in an open field test apparatus with a luminance of 70 lx. The test consisted of a habituation session (the first test session) and the second test session. Each session lasted 15 min and occurred in the following order. In the habituation session, two empty cylindrical wire cages (inner size, 7 cm φ × 15 cm [H]; outer size, 9 cm φ × 16.5 cm [H], with 21 vertical stainless (3-mm-φ wires) longitudinally and gray polyvinyl discs on the top and bottom, manufactured by the RIKEN Rapid Engineering Team) were placed in two adjacent corners. In the first test session, a mouse (7-week-old male DBA2, purchased from Nihon SLC, Shizuoka, Japan) that was new to the test mouse was placed in one of two cylindrical cages. In the second test session, another mouse that was also new to the test mouse was placed in the remaining cylindrical cage. Between the three sessions, there were 4-min intervals during which the test mouse was returned to its home cage. The three sessions were video-recorded from above, and the times spent in the two corner squares containing the cylinders within the 3 × 3-square subdivision (17.7 × 17.7 $cm^2$) were measured with Image J OF4 (O'Hara). For the two test sessions, video recording was also performed from an obliquely upward position to observe contact between the test mouse and the in-cage mouse. Contact with the in-cage mouse was defined as a forward movement toward the mouse in the cage and subsequent direct contact with the head. The position and posture of the in-cage mice were observed through the slits of the wires. The contacts were counted on the video records by an observer who was blind to the genotypes. Each in-cage mouse was used once a day, and when the habituation session began, the mouse was simultaneously placed in its cylindrical cage on the corners of an open field box that was not being used for the tests. These rules were thought to minimize the difference between the two in-cage mice in the second test session concerning their acclimatization to the cylindrical cage and the open field box environment. After each use, the cylindrical cage was extensively washed with water and rinsed with 90% ethanol, which was then evaporated to minimize the effects of the remaining materials.

*Tail suspension test*

The tail suspension test was conducted as previously described (36). Mice were attached to a wire using an adhesive tape placed approximately 1.5 cm from the tip of the tail and suspended 30 cm above the floor. The duration of immobility was recorded for 5 min.

*Marble burying test*

Transparent plastic cages (25×18×15 cm) were filled with white paper bedding material (Paperclean, Japan SLC) to a 4-cm depth. Mice were placed individually in the test cages for 20 min (habituation trial) and then returned to their home cages. Fifteen blue glass marbles were evenly set at 5-cm intervals on the surface of the bedding material in the habituated cages. Then the mice were again placed in the habituated test cage for 20 min (test trial). During both trials, the test cage with the mouse was set in the infrared beam apparatus to measure activity. After 20 min, marbles that were more than two-thirds covered with paper clean were counted as buried marble.

*Classical fear conditioning test*

This test consisted of three parts: a conditioning trial (Day 1), a context test trial (Day 2), and a cued test trial (Day 3). Fear conditioning was performed in a clear plastic chamber equipped with a stainless-steel grid floor [34626630 (H) cm]. A CCD camera was mounted on the ceiling of the chamber and connected to a video monitor and computer. The grid floor was wired to a shock generator. White noise (65 dB) was supplied from a loudspeaker as an auditory cue [i.e. the conditioned stimulus (CS)]. The conditioning trial consisted of a 2-min exploration period followed by two CS–US pairings separated by 1 min. A US (foot-shock:0.5 mA, 2 s) was administered at the end of the 30-s CS period. Twenty-four hours after the conditioning trial, a context test was performed in the same conditioning chamber for 3 min in the absence of the white noise. A cued test was also performed in an alternative context with distinct cues; the test chamber was different from the conditioning chamber in terms of luminance (about 0 to 1lx), color (white), floor structure [no grid but with thin bedding material (Alpha-Dri: Shepherd, TN, USA)], and shape (triangular).The cued test was conducted 24 h after the contextual test was finished; it consisted of a 2-min exploration period (no CS) to evaluate nonspecific contextual fear, followed by a 2-min CS period(no foot shock) to evaluate the acquired cued fear. The rate of freezing response (immobility, except for respiration and heartbeat) of mice was measured as an index of fear memory. Data were collected and analyzed with Image J FZ2 (O'Hara).

*Acoustic startle response and prepulse inhibition*

Mice were habituated in their home cages for 1 h to 65-dB white noise. They were then placed into standard startle chambers. Each session was initiated with a 5-min acclimation period of white noise at 65 dB followed by 10 successive 120-dB tones to elicit the startle response (40 ms). Nine different trial types were then presented: 70, 75, 80, 85, 90,95, 100, 110 or 120 dB (40 ms) with background noise at 65 dB. Each trial was presented five times, and the average response to each trial calculated. Immediately after the startle response trials, the PPI session was begun. During each PPI session, a mouse was exposed to the following types of trials: the omission of stimuli (no-stimulus trial); startle-alone trial (120 dB); three prepulse combinations (prepulse-pulse trial) using three prepulse intensities: 70, 75 and 80 dB. Each PPI session consisted of 10 presentations of each type of trial. PPI was assessed for each animal as a percentage (%PPI): 100(mean startle to pulse alone—mean startle to prepulse-pulse)/mean startle to pulse alone. The apparatuses and software used for data analysis were commercially available (Mouse Startle; O'Hara).

*Protein Modeling of WT and mutant ST3*

Based on the P-score 0.2, top model was selected. For further refinement of the homology model, Galaxyrefine server (https://galaxy.seoklab.org/) was used to improve the overall quality of the model. The constructed homology model was validated using the Ramachandran plot server (zlab.umassmed.ed) to assist with the quality of the model. The model shows 89.3% to be in the allowed region, 9.1% is prefer regions, while only 1.6 percent of the residues showed questionable. The overall quality of the homology model was validated using the ProSA-web server (https://prosa.services.came.sbg.ac.at) to assess the protein structure quality. The quality score of the homology model is (Z-Score: -8.26) is in the accepted range. The homology wild type model (WT) was selected in Chimera to mutate the desired residues and the mutated structure was minimized by 1000 steps of steepest decent algorithm followed by 5000 steps of conjugate gradient minimization. The superposition of the wild type and mutated models ensures the calculation of the RMSD using Chimera v.1.16. The WT (cyan) and mutated (red) structures were aligned and represented in cartoon representation emphasizing the mutated residues (WT-CYS: yellow, Mutated-ARG: green) in a ball and stick representation of the ST3 protein. The root mean square deviation (RMSD) was calculated (248 pruned atoms; 1.433 angstroms) to analyze the changes in structure and dynamics of the modelled proteins.

#### Supplementary figures

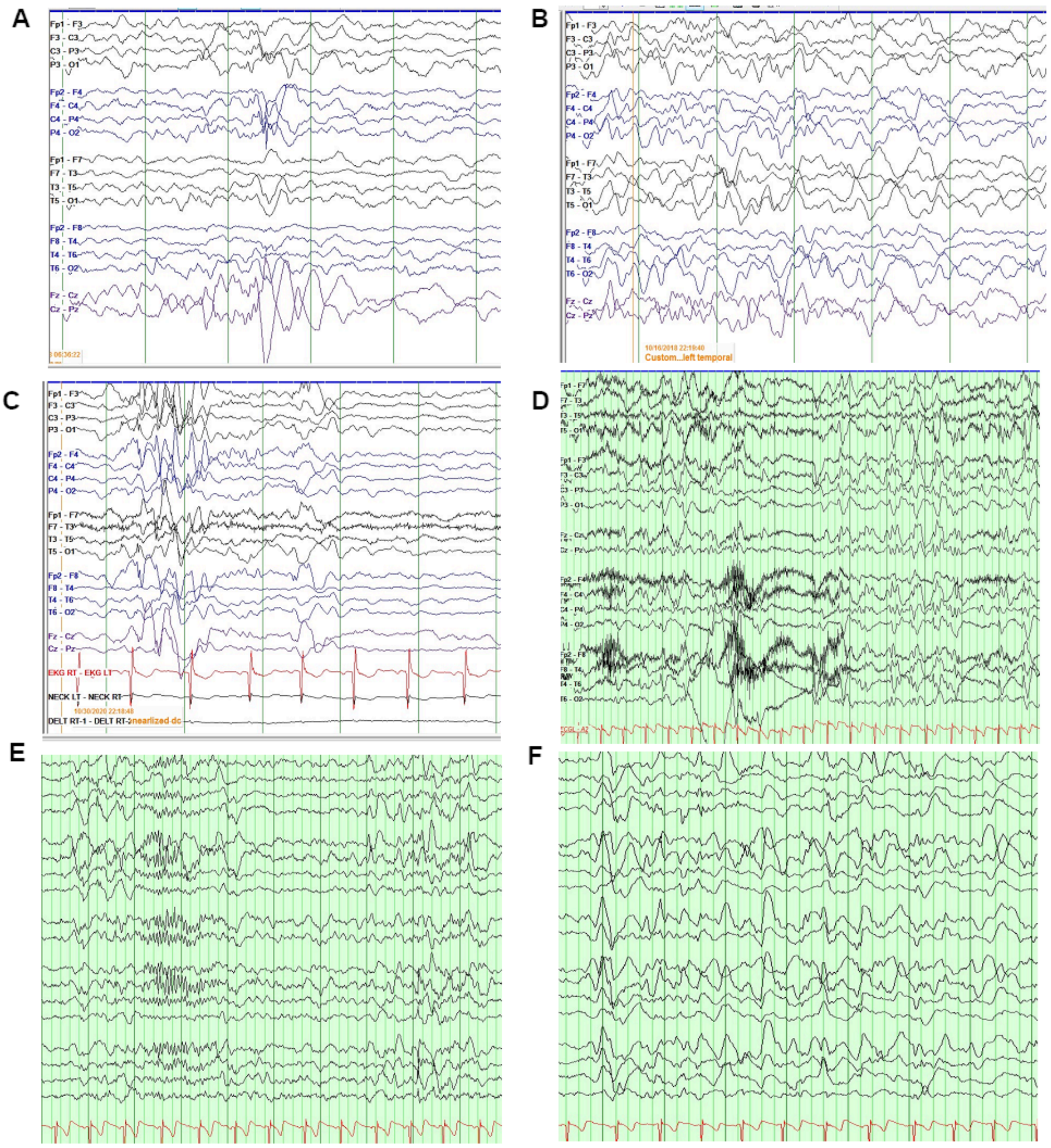


**Supplementary Fig 1. EEG studies.** Sleep EEG at last follow-up of patient 5 showing slowed background, right (A) and left (B) central and anterior slow and sharp waves; diffuse epileptic discharges (C). Awake EEG at last follow-up of patient 6 showing diffuse slow spike slow waves and irregular slow spike-and-slow wave complexes within poorly organized and slowed background activity (D). Sleep EEG showing generalized paroxysmal fast activity prominent over the anterior regions (E) and intermittent generalized delta activity (F).

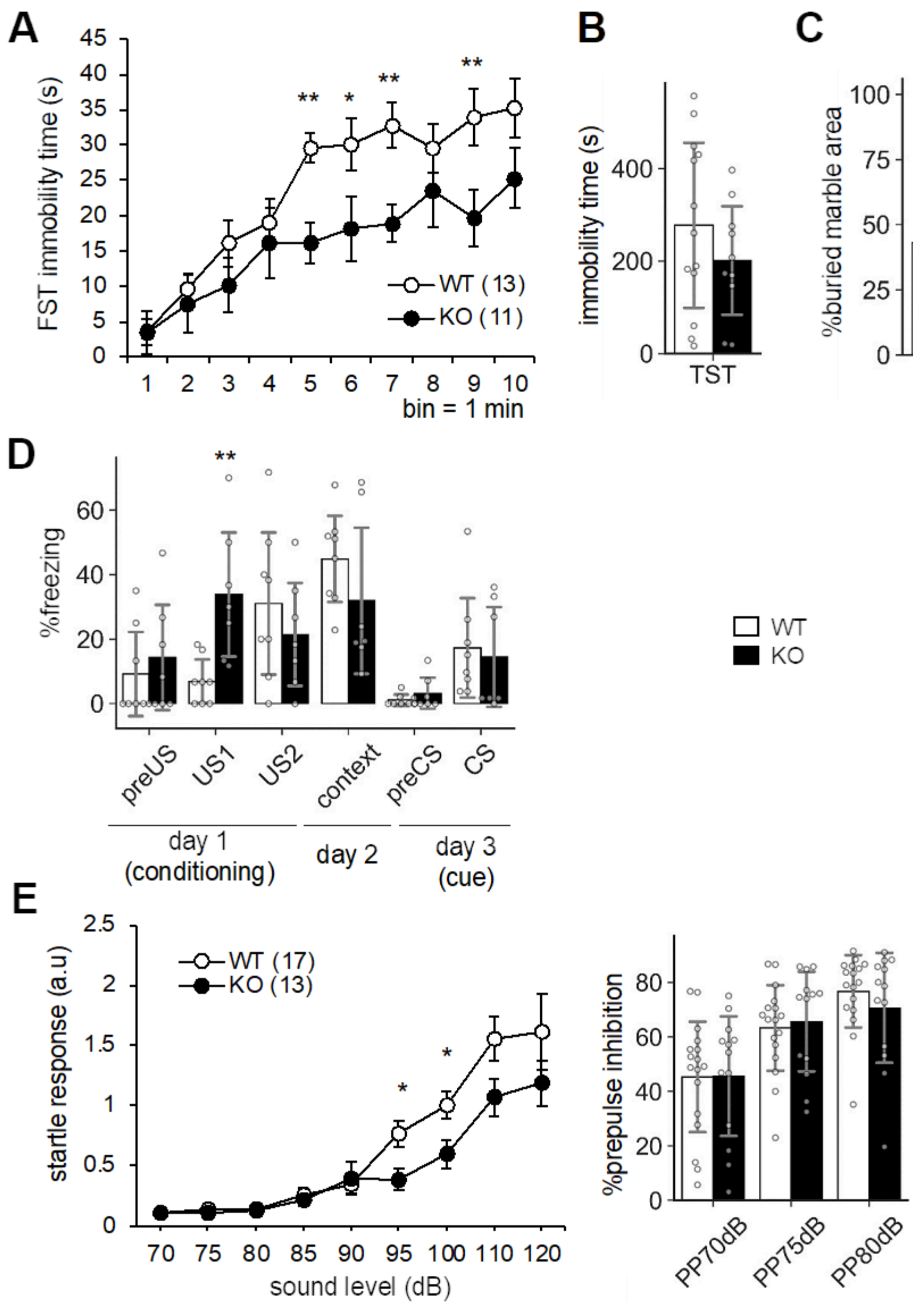


**Supplementary Fig 2. Supplementary results for ST3 KO behavioral analysis.** (A) Time course of immobility in forced swimming test. (B) Immobile time in tail suspension test (10 min). (C) Marble burying test. Percentages of the buried marble area was quantified. (D) Fear conditioning test. (E) Startle response (left) and its prepulse inhibition (right). Mean values are indicated in all graphs. *Open bars and circles*, WT; *closed bars and circles*, KO. *Error bars*, SD (bar graphs), SEM (line graphs). *Gray circles on bar graphs* indicate the individual value of each mouse. The *numbers in parentheses* in line graphs indicate n (the number of mice) in each experimental group. *, $p < 0.05$; **, $p < 0.01$; ***, $p < 0.001$ in two-tailed unpaired Student's t-test.

A

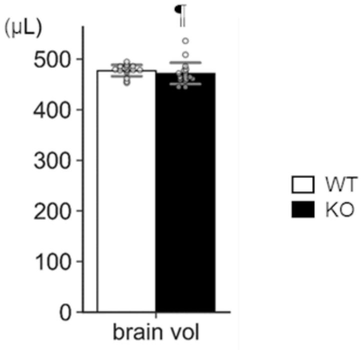


B

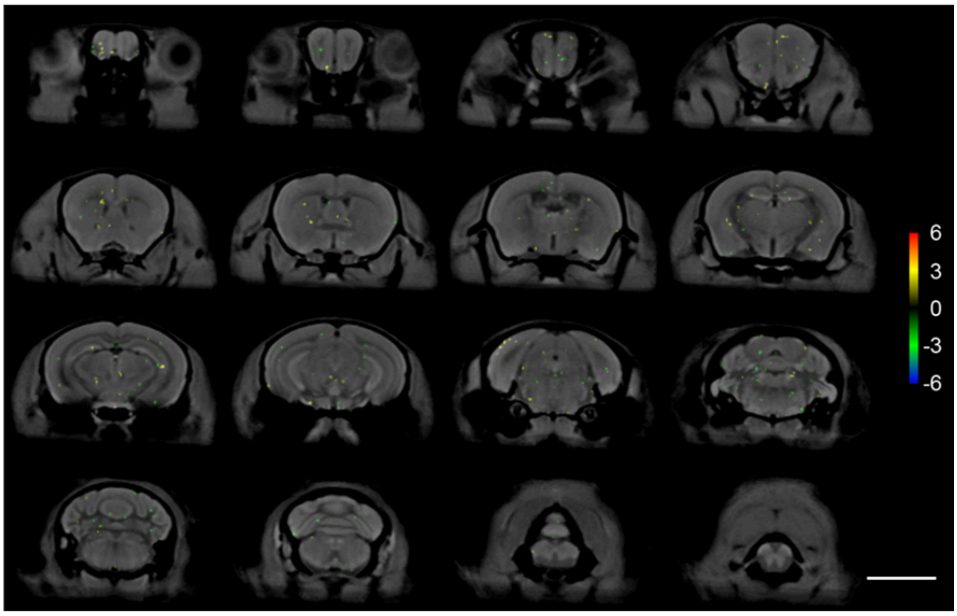


**Supplementary Fig 3. MRI analysis of ST3 KO brains**. (A) Total brain volumes. Mean values are indicated. Open bars and circles, WT; closed bars and circles, KO. Error bars, SD. Gray circles indicate the individual value of each mouse. ¶ $p < 0.05$ in F-test. There were no significant difference in two-tailed unpaired Welch's t-test. WT, n = 16 mice; KO, n = 18 mice. (B) Results of tensor-based morphometric analysis. t-statistic map overlaid upon the mean of all registered images. The clusters are small and sporadically distributed. All of the signals disappear after any sort of control for multiple comparisons (Bonferroni, Holm-Bonferroni, or False Discovery Rate). Scale bar, 5 mm. WT, n = 16 mice; KO, n = 18 mice.

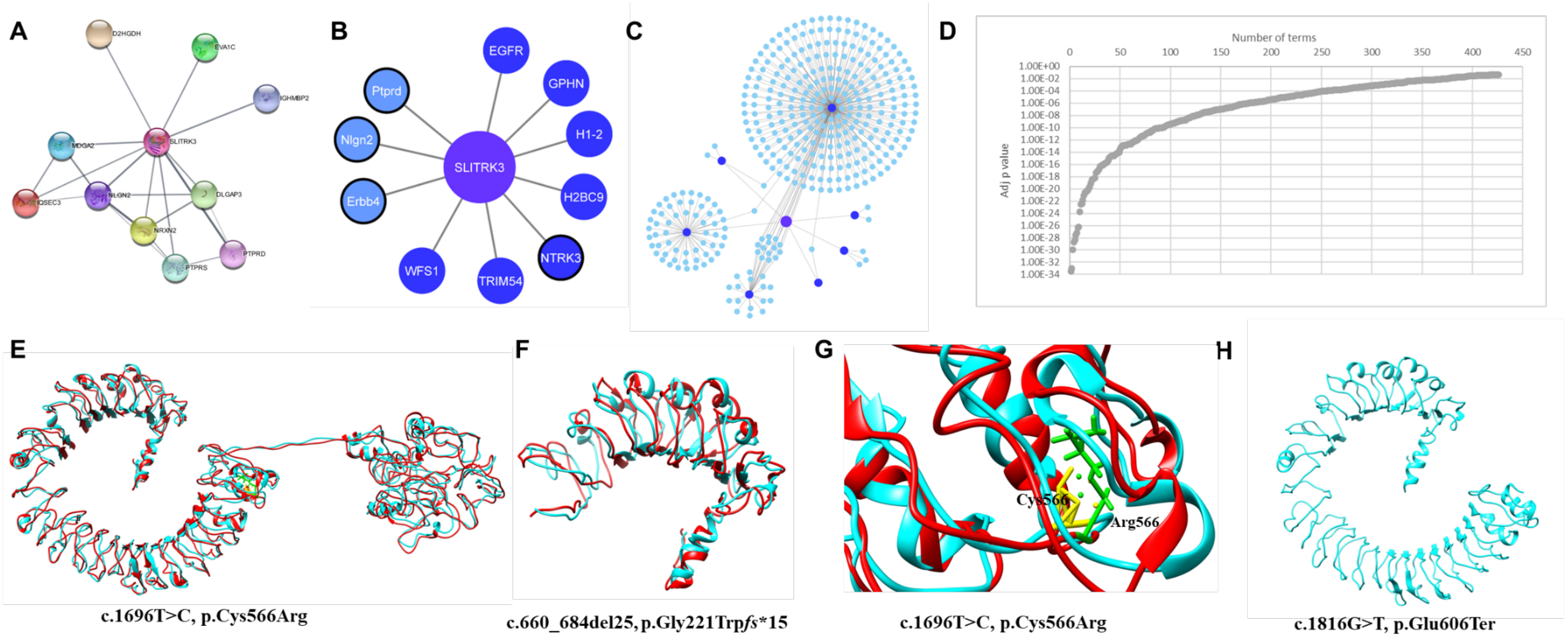


**Supplementary Fig 4. *In-silico* network analysis and structural modelling of wildtype and mutant SLITRK3 proteins**. (A) General protein-protein network showing different interactions of candidate genes with SLITRK3. (B) Direct interactors of SLITRK3 (1$^{st}$ layer interactome). Black border indicates that they resulted from the manual curation of the literature. The colour of the interactors is coded based on the confidence of the result. Blue: lower confidence, Darker blue: higher confidence. (C) Direct and indirect interactors of SLITRK3 (2$^{nd}$ layer interactome). The interactors of the high confidence direct interactors of SLITRK3 are presented and coloured in light blue. The order of the direct interactors (darker bluer) is alphabetical and starts from the top in a clockwise manner: EGFR, GPHN, H1-2, H2BC9, NTRK3, TRIM54, and WFS1. B&C were produced using Cytoscape v3.9.0. (D) Selecting a threshold for the GOBP enrichment. The distribution of adjusted p values of the enrichment of biological processes of the 2$^{nd}$ layer interactome of SLITRK3 is plotted, as resulted from g:Profiler. The red line indicates the selected threshold. (E) Predicted tertiary structure and homology model of the wild type SLITRK3 protein using I-TASSER Structure Prediction server as well as of (F) the mutated c.1696T>C, p.Cys566Arg variant with a disrupted disulphide bridge resulting in altered protein structure confirmation, and the (G) nonsense c.1816G>T, p.Glu606Ter variant showing a shorter protein resulting in the termination of the chain and loss of the cytoplasmic domain of the protein. (H) Structural modelling of the frameshift variant c.660_684del25, p.Gly221Trp*fs**15 in SLITRK3 using Chimera v.1.4, in comparison with the wild-type SLITRK3 protein showing the presence of only a chunk of the extracellular domain being formed.

### Supplementary Table 1. Animal list.

| Supplementary Table 1. Animal list. | | | | | | | | | |
|---|---|---|---|---|---|---|---|---|---|
| related displaying items | subjects | cage# | sex | age (weeks-old) | genotype# | | | total mice# | excluded mice# |
| | | | | | WT | HET | KO | | |
| Fig. 3A | survival curves for B6N8 mice | 6 | M, F | 0 - 7 | 7 | 24 | 9 | 40 | 0 |
| Fig. 3B | body weight for B6N8 mice | 7 | M, F | 2 - 3 | 11 | 23 | 10 | 44 | 0 |
| Fig. 3B | body weight for B6N1/2 mice | 5 | M | 9 - 14 | 10 | 0 | 9 | 19 | 0 |
| Fig. 4A | homecage activity | 3 | M | 9 - 14 | 11 | 0 | 10 | 21 | 1 (died) |
| Fig. 4B | rotarod test | 7 | M | 26 - 27 | 13 | 0 | 11 | 24 | 0 |
| Fig. 4C | elevated plus maze test for 5 weeks-old mice | 3 | M | 5 | 8 | 0 | 7 | 15 | 0 |
| Fig. 4C | elevated plus maze test for 12 weeks-old mice | 3 | M | 12 | 11 | 0 | 9 | 20 | 0 |
| Fig. 4D | forced swimming test | 7 | M | 28 - 31 | 13 | 0 | 11 | 24 | 0 |
| Fig. 4E | wire hanging test | 4 | M | 25 - 27 | 5 | 0 | 4 | 9 | 0 |
| Fig. 4F | novel object approach test | 3 | M | 10 - 13 | 11 | 0 | 9 | 20 | 0 |
| Fig. 4G | resident intruder test | 6 | M | 13 - 15 | 8 | 0 | 7 | 15 | 0 |
| Fig. 4H | social interaction test | 6 | M | 12 - 14 | 14 | 0 | 12 | 26 | 0 |
| Fig. 5 | interneuron counting | 4 | M, F | 2 | 7 | 0 | 7 | 14 | 0 |
| Table 3 | lethality for B6N1/N2 mice | 123 | M, F | 4 - 7 | 280 | 527 | 166 | 973 | 0 |
| Table 3 | lethality for B6N8 mice | 29 | M, F | 4 - 7 | 64 | 93 | 17 | 174 | 0 |
| Suppl Fig. 2A | forced swimming test | 7 | M | 28 - 31 | 13 | 0 | 11 | 24 | 0 |
| Suppl Fig. 2B | tail suspension test | 8 | M | 30 | 13 | 0 | 10 | 23 | 0 |
| Suppl Fig. 2C | marble burying test | 9 | M | 20 - 24 | 13 | 0 | 12 | 25 | 0 |
| Suppl Fig. 2D | fear conditioning test | 5 | M | 14 - 20 | 8 | 0 | 7 | 15 | 0 |
| Suppl Fig. 2E | auditory startle response/prepuse inhibition | 12 | M | 12 - 19 | 17 | 0 | 13 | 30 | 0 |
| Suppl Fig. 3 | magnetic resonance imaging | 14 | M | 17 - 28 | 16 | 0 | 18 | 34 | 0 |

#### Supplementary Table 2. Top enriched pathways based on the adjusted p value.

| Pathway | Term id | Adjusted p value | Intersection size |
|---|---|---|---|
| Signaling by Receptor Tyrosine Kinases | REAC:R-HSA-9006934 | 3.91E-35 | 79 |
| Signal Transduction | REAC:R-HSA-162582 | 1.34E-20 | 150 |
| Immune System | REAC:R-HSA-168256 | 7.26E-20 | 131 |
| Signaling by Interleukins | REAC:R-HSA-449147 | 4.78E-18 | 56 |
| Cytokine Signaling in Immune system | REAC:R-HSA-1280215 | 1.50E-17 | 69 |
| Signaling by EGFR | REAC:R-HSA-177929 | 2.05E-17 | 21 |
| Signaling by ERBB2 | REAC:R-HSA-1227986 | 5.43E-15 | 19 |
| Diseases of signal transduction by growth factor receptors and second messengers | REAC:R-HSA-5663202 | 4.66E-14 | 48 |
| Axon guidance | REAC:R-HSA-422475 | 9.44E-14 | 55 |
| Signaling by SCF-KIT | REAC:R-HSA-1433557 | 1.07E-13 | 17 |
| Disease | REAC:R-HSA-1643685 | 2.31E-13 | 105 |
| Nervous system development | REAC:R-HSA-9675108 | 6.79E-13 | 55 |
| VEGFA-VEGFR2 Pathway | REAC:R-HSA-4420097 | 1.03E-12 | 23 |
| Downstream signal transduction | REAC:R-HSA-186763 | 1.34E-12 | 14 |
| Signaling by ERBB2 in Cancer | REAC:R-HSA-1227990 | 2.75E-12 | 13 |
| Clathrin-mediated endocytosis | REAC:R-HSA-8856828 | 3.87E-12 | 27 |
| Interleukin-3, Interleukin-5 and GM-CSF signaling | REAC:R-HSA-512988 | 4.67E-12 | 17 |

#### Supplementary Table 3. Top enriched pathways based on the number of proteins of the SLITRK3.

| Pathway name | Term id | Adjusted p value | Intersection size |
|---|---|---|---|
| Signal Transduction | REAC:R-HSA-162582 | 1.34E-20 | 150 |
| Immune System | REAC:R-HSA-168256 | 7.26E-20 | 131 |
| Disease | REAC:R-HSA-1643685 | 2.31E-13 | 105 |
| Signaling by Receptor Tyrosine Kinases | REAC:R-HSA-9006934 | 3.91E-35 | 79 |
| Cytokine Signaling in Immune system | REAC:R-HSA-1280215 | 1.50E-17 | 69 |
| Developmental Biology | REAC:R-HSA-1266738 | 2.17E-07 | 69 |
| Innate Immune System | REAC:R-HSA-168249 | 6.24E-07 | 67 |
| Infectious disease | REAC:R-HSA-5663205 | 2.07E-08 | 63 |
